# Hallucinating protein assemblies

**DOI:** 10.1101/2022.06.09.493773

**Authors:** B. I. M. Wicky, L. F. Milles, A. Courbet, R. J. Ragotte, J. Dauparas, E. Kinfu, S. Tipps, R. D. Kibler, M. Baek, F. DiMaio, X. Li, L. Carter, A. Kang, H. Nguyen, A. K. Bera, D. Baker

**Author notes:** These authors contributed equally to this work (B. I. M. Wicky, L. F. Milles, A. Courbet).

## Abstract

Deep learning generative approaches provide an opportunity to broadly explore protein structure space beyond the sequences and structures of natural proteins. Here we use deep network hallucination to generate a wide range of symmetric protein homo-oligomers given only a specification of the number of protomers and the protomer length. Crystal structures of 7 designs are very close to the computational models (median RMSD: 0.6 Å), as are 3 cryoEM structures of giant rings with up to 1550 residues, C33 symmetry, and 10 nanometer in diameter; all differ considerably from previously solved structures. Our results highlight the rich diversity of new protein structures that can be created using deep learning, and pave the way for the design of increasingly complex nanomachines and biomaterials.

**One-sentence summary:** Deep network-based protein design enables the generation of cyclic homo-oligomers across the nanoscopic scale.

## Main text

Cyclic protein oligomers play key roles in almost all biological processes and constitute nearly 30% of all deposited structures in the Protein Data Bank (PDB, (*1*) (*2–4*)). Because of the many applications of cyclic protein oligomers, ranging from small molecule binding and catalysis to building blocks for nanocage assemblies(*5*), *de novo* design of such structures has been of considerable interest from the beginning of the protein design field (*6, 7*). While there have been a number of successes (*8–10*), current approaches require specification of the structure of the protomers in advance, and with the exception of parametrically designed helical bundles (*11, 12*), have involved rigid body docking of previously characterized monomers into higher order symmetric structures followed by interface optimization to confer low energy to the assembled state (*13–17*). The requirement that the protomer structure be specified in advance has limited exploration of the full space of oligomeric structures; in particular assemblies in which the chains are more intertwined. For monomeric proteins, broad exploration of the space of possible structures has become possible by deep network hallucination: starting from a random amino acid sequence, Monte Carlo optimization with a loss function favoring folding to a well-defined state converges on new sequences that fold to new structures (*18–21*). We reasoned that deep network hallucination could enable the design of higher-order protein assemblies in one step, without prespecification or experimental confirmation of the structures of the protomers, provided that a suitable loss function could be formulated (*18–20, 22–24*).

We set out to broadly explore the space of cyclic protein homo-oligomers by developing a method for hallucinating such structures that places no constraints on the structures of either the protomers or the overall assemblies. Starting from only a choice of chain length *L* and oligomer valency *N* (2 for a dimer, 3 for a trimer, etc.), the method initializes a random amino acid sequence to begin a Monte Carlo search in sequence space (Fig. 1A). The loss function guiding the search is computed by inputting *N* copies of the sequence into the AlphaFold2 (AF2) network (*25*), and combining structure prediction confidence metrics (pLDDT and pTM) with a measure of cyclic symmetry; the standard deviation of the distances between the center of mass of adjacent protomers within the predicted structure.

**Fig. 1.**
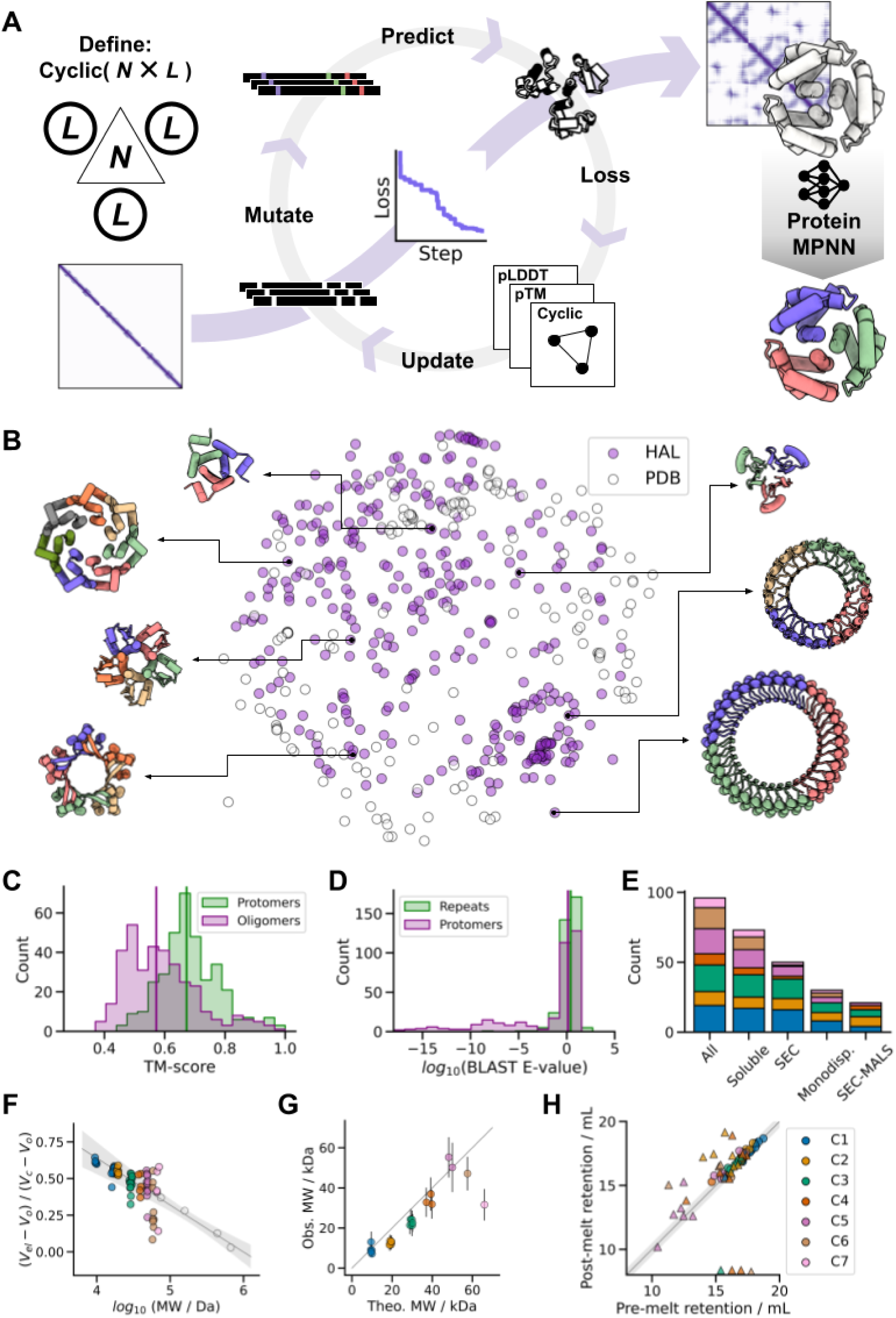
Hallucinating protein assemblies. (**A**) Starting from the definition of a cyclic symmetry and protein length, a random sequence is optimized by MCMC through the AF2 network until the resulting structure fits the design objective, followed by sequence re-design with ProteinMPNN. (**B**) The method generates structurally diverse outputs, quantified here by multi-dimensional scaling of protomer pairwise structural similarities between experimentally tested HALs (*N* = 351) and all *de novo* cyclic oligomers present in the PDB (*N* = 162). (**C**) Generated structures are significantly different from anything present in the PDB. Median TM-scores to the closest match: 0.67 and 0.57 for the protomers and oligomers respectively (vertical lines). (**D**) Generated sequences are unrelated to naturally-occuring proteins. Median BLAST E-values from the closet hit in UniRef100: 2.6 and 1.3 for the repeat motifs and protomers respectively (vertical lines). (**E**) Success counts of ProteinMPNN-designed HALs at different levels of characterization. (**F**) Most soluble HALs have SEC retention volumes consistent with their oligomeric state. The gray line shows the fit to calibration standards (open circles), and the shaded area represents the 95% confidence interval of the calibration. (**G**) Parity plot between the theoretical and observed molecular weights of HALs from SEC-MALS. (**H**) ProteinMPNN-designed HALs are thermostable. Parity plot between pre-melting and post-melting retention volumes; circles represent designs that remained monodisperse, while triangles indicate polydispersity after heating the sample. In plots **E**-**H,** the data is categorized by symmetries. The legend is shown in **H**.

We found that monomers and dimeric to heptameric assemblies could readily be generated by this procedure for chains of 65 to 130 amino acids, with converging trajectories typically coalescing to cyclic homo-oligomeric structures within a few hundred steps (approximately one week of CPU-time). The resulting structures are topologically diverse, spanning all-α, mixed α/β and all-β structures and differ from structurally-verified cyclic *de novo* designs present in the PDB (Fig. 1B). These assemblies, which we term HALs, also differ from natural proteins, with the median closest relatives in the PDB having TM-scores of 0.67 and 0.57 for the protomers and oligomers respectively (29% of the structures have TM-scores < 0.5, which constitutes the cutoff for fold assignment in CATH/SCOP (*26*) (Fig. 1C), and sequences unrelated to natural ones (Fig. 1D), indicating considerable generalization beyond the PDB training set.

We selected 150 designs with pLDDT > 0.7 and pTM > 0.7 for experimental testing. However, virtually none showed significant soluble expression when produced in *E. coli* (median soluble yield: 9 mg per liter of culture-equivalent, Fig. S1), and of the few that were marginally soluble none had both the expected oligomerization state by size-exclusion chromatography (SEC) and a circular dichroism (CD) profile consistent with the hallucinated structure. We speculated that this failure could be a consequence of over-fitting during MCMC optimization leading to the generation of adversarial sequences (see Fig. S2). Analogous neural network activation maximization approaches with 2D images similarly can lead to non-viable solutions (*27–29*). To eliminate such over-fitting, we generated new sequences for the hallucinated oligomer backbones using the recently developed ProteinMPNN sequence design method (*Dauparas et al.: https://www.biorxiv.org/content/10.1101/2022.06.03.494563v1*). For each original backbone, 24 to 48 sequences were generated with ProteinMPNN, and assembly to the target oligomeric structure validated with AF2 (these evaluations are far fewer in number compared to the thousands of evaluations in the original hallucination trajectories, making overfitting much less likely). We independently evaluated the designs using an updated version of RoseTTAFold (RF2) (*30*) and found that while most of the original AF2 hallucinated sequences were not confidently predicted to fold to the hallucinated structures (see Fig. S2, S6, S7), following ProteinMPNN redesign almost all were predicted to fold correctly.

We tested 96 ProteinMPNN-designed HALs with pLDDT > 0.75 and RMSD to original backbone < 1.5 Å and found that 71/96 (74%) showed of high levels of soluble expression (median yield: 247 mg per liter of culture-equivalent), 50/96 (52%) had a SEC retention volume consistent with the oligomeric size (of which 30 (60%) were monodisperse) (Fig. 1F and Fig. S3), and at least 21/96 (22%) had the correct oligomeric state when assessed by SEC-Multi Angle Light Scattering (SEC-MALS) (Fig. 1G). Furthermore, CD analysis of the soluble samples indicated that 67/71 (96%) had secondary structure contents consistent with the designs (Fig. S4). These success rates are in stark contrast to those of the original AF2 sequences, indicating that the MCMC hallucination procedure generates viable backbones, but over-fitted sequences, and highlighting the power of ProteinMPNN to generate sequences which fold to a given backbone structure (Fig. 1E). We assessed the thermal stability of the 71 soluble HALs by CD spectroscopy, and found that 54 maintained their secondary structure up to 95 °C (Fig. S4). SEC characterization of the heated-treated samples indicated that most designs retained their oligomeric state, suggesting that the HAL assemblies are thermostable (Fig. 1H, S4).

To evaluate design accuracy we attempted crystallization of 19 designs and succeeded in solving crystal structures for seven (three C2s, two C3s and two C4s) (Fig. 2). All crystal structures had the correct oligomerization state and closely matched the design models (median Cα RMSD of 0.6 Å across all designs, Fig. 2 and Fig. S5). The side chain conformations in the crystal structures also closely match those in the design models (Fig. 2).

**Fig. 2.**
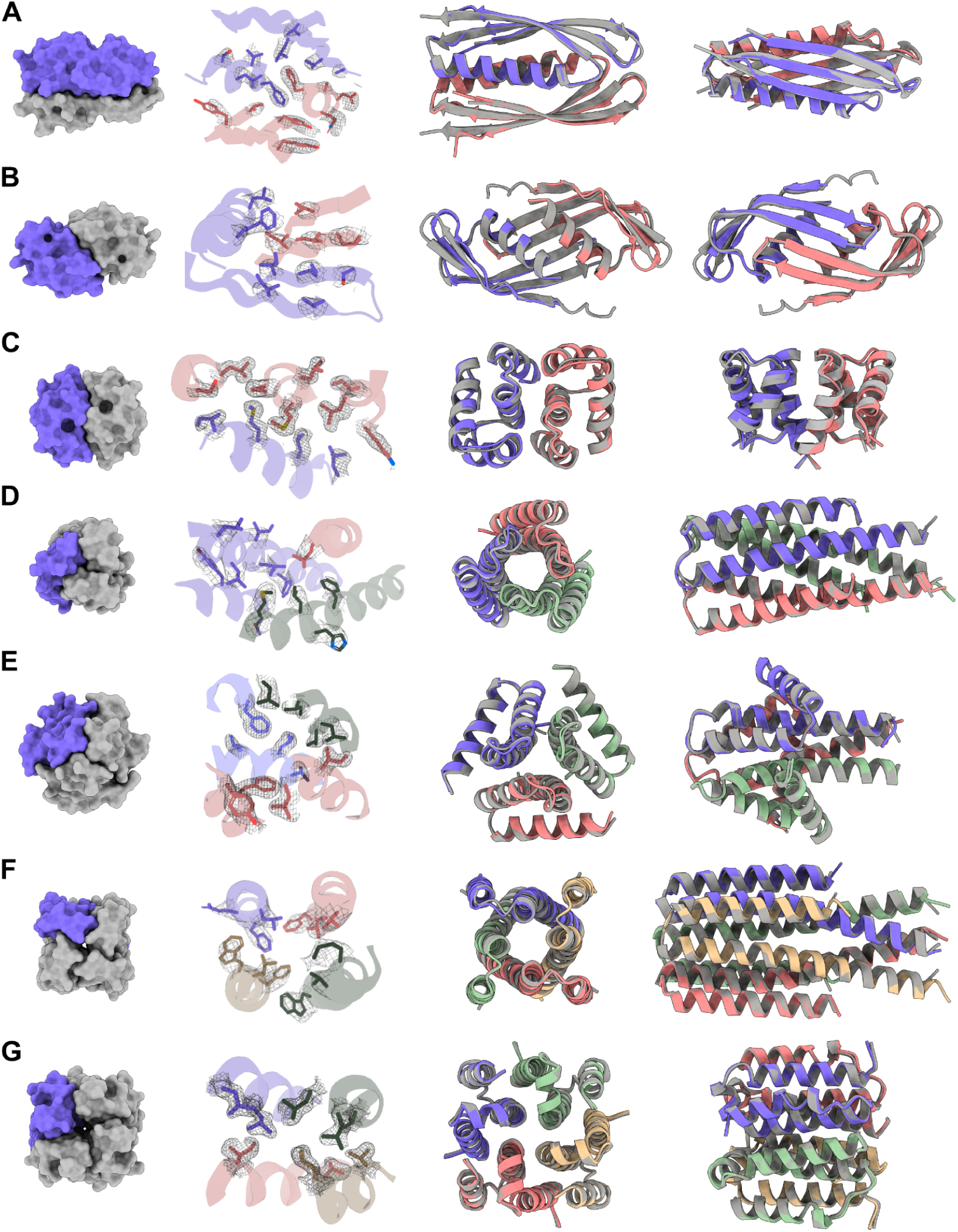
Structures of HALs solved by X-ray crystallography compared to their design models. (**A**) HALC2_062 (RMSD: 0.81 Å). (**B**) HALC2_065 (RMSD: 1.02 Å). (**C**) HALC2_068 (RMSD: 0.86 Å). (**D**) HALC3_104 (RMSD: 0.42 Å). (**E**) HALC3_109 (RMSD: 0.46 Å). (**F**) HALC4_135 (RMSD: 0.60 Å). (**G**) HALC4_136 (RMSD: 0.34 Å). In each row, the first panel shows a surface rendering of the oligomer with one protomer highlighted in purple, the second, the 2mFo-DFc map in gray compared to the side-chain rotamers of the design model, and the last two panels, two different orientations of the structural overlays between the model (gray) and the solved structure (color by chains).

The solved structures exhibit striking diversity with many intricate structural features. HALC2_062 (Fig. 2A) is a three-layer homo-dimer with a single helix from each protomer packed together between two outer β-sheets (one from each protomer), while HALC2_065 (Fig. 2B) is also a mixed α/β homo-dimer, but has a single, continuous β-sheet shared between both chains, which wraps around two perpendicular paired helices. These two hallucinated structures are very different from anything deposited in the PDB, with TM-scores to their best matches of 0.59 and 0.54 respectively (Fig. 4A-B, Table S1). HALC2_068 (Fig. 2C) is a fully helical dimer with an extensive interface formed by 6 interacting helices (3 from each protomer), with a single perpendicular helix buttressing the interfacial helices. Despite the absence of secondary structure complexity and long-range contacts, this design also differs significantly from its closest structural relative in the PDB (TM-score: 0.57, Fig. 4C, Table S1). HALC3_104 (Fig. 2D) is a homo-trimeric coiled-coil, with a central bundle of three helices, augmented by an outer-ring of three shorter helices that lay in the groove formed by adjacent protomers. Unsurprisingly given the simplicity of this topology, there is a close structural match in the PDB (TM-score: 0.88, Fig. 4D, Table S1). HALC3_109 (Fig. 2E) is a homo-trimeric three-layer all-helical structure, with three inner helices splaying outwards to contact two additional helices from the same protomers at angles of roughly 25° and 90°; the closest assembly in the PDB has a TM-score of 0.69 (Fig. 4E, Table S1). HALC4_135 (Fig. 2F) is a coiled-coil composed of helical hairpins reminiscent of HALC3_104, but with C4 symmetry instead of C3, and a discontinuous superhelical twist. Despite its simple topology, the closest structural homologue of this design has a TM-score of only 0.59 (Fig. 4F, Table S1). HALC4_136 (Fig. 2G) is composed of 3-helix protomers with eight outer helices encasing four almost fully hydrophobic inner helices, where two of the helices are rigidly linked through a 90° helical kink. The closest match in the PDB has a TM-score of 0.71, but the matched structure has C5 symmetry rather than the C4 symmetry of the design and crystal structure.

Next, we sought to generate HALs of increased complexity across longer length-scales by extending the design specifications to structures of higher symmetry (up to C42) and longer assembly sequence length (up to 1800 residues). To generate multiple possible oligomers from a single structure, we specified the MCMC trajectories as single-chains with internal sequence symmetry, with the goal of generating structure-symmetric repeat proteins that could be split into any desired oligomeric assembly compatible with factorization (e.g. C15 into a pentamer, shorthanded as C15-5). To maximize the exploration of the design space while minimizing use of computational resources, we devised an evolution-based computational strategy: many short MCMC trajectories (< 50 steps) outputs were clustered by structure prediction confidence metrics (pLDDT and pTM), and then used to seed new trajectories (see Supplementary Materials). Using this approach, we hallucinated cyclic homo-oligomers from C5 to C42 ranging from 7 to 14 nm (median: 10 nm) along their largest dimension, which were then divided into homo-trimers, tetramers, pentamers, hexamers, heptamers, octamers, and dodecamer, and the backbones were re-designed with ProteinMPNN (Fig 1C). While the α/β topology of some of these larger HALs is reminiscent of natural Leucine Rich Repeats (LRRs, (*31*)), which is reflected by a median highest protomer TM-scores of 0.64, these ring-shaped structures differ considerably from the horseshoe folds of LRRs that do not close into cyclic structures. The closest oligomer structures in the PDB have a median TM-score of 0.47, and BLAST sequence similarity searches for the repetitive sequence motif do not return any significant hits (Fig. 1D); the hallucination process as in the earlier cases clearly generalizes well beyond the training set.

These larger HALs have overall molecular weights greater than 100 kDa, and thus were well-suited for structural characterization by electron microscopy (EM). We subjected soluble large HALs with a SEC retention volume consistent with the size of their oligomeric state to screening by negative stain EM (nsEM). Inspecting the resulting micrographs, we found that all of the designs screened showed monodisperse particles of the expected size and circular shape (Fig. S8). We obtained 2D class averages and 3D *ab initio* reconstructed electron density maps for six designs (two C5s, three C6s, and one C7) with C6 to C42 internal repeat symmetry that clearly showed low-resolution structural features and diameters consistent with their designs (Fig. 3A, Fig. S12). Next, we selected three designs: one C15 homo-pentamer (HALC5-15_262), one C18 homo-hexamer (HALC6-18_265) and one C33 homo-trimer (HALC3-33_343) for high-resolution single particle cryoEM characterization. We collected datasets that produced 2D class averages with clear secondary structure feature placements, and 3D *ab initio* reconstruction and refinement yielded 3D electron density maps at 4.38 Å, 6.51 Å and 6.32 Å resolution respectively. HALC5-15_262 was designed as a homo-hexamer, but structure prediction calculations were more consistent with a pentameric structure with a nearly identical protomer internal conformation and a very slightly shifted subunit interface (Fig. S13); the cryoEM structure is also a pentamer with an Cα RMSD of 1.69 Å to the predicted structure.

**Fig. 3.**
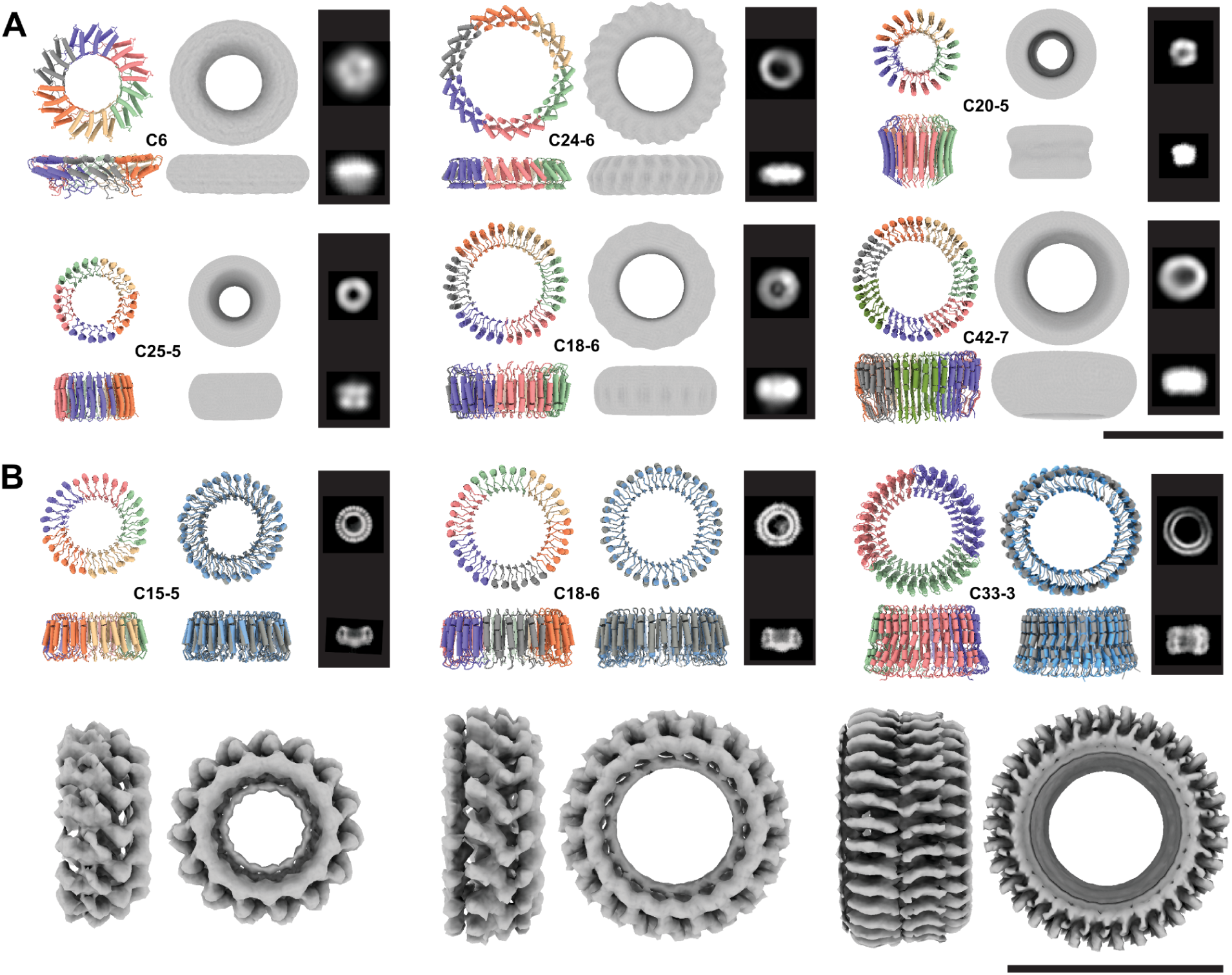
Cryo-electron and negative stain electron microscopy validation of large HALs. For each design, the model is shown colored by chain and the corresponding internal symmetry (X) and oligomerization state (Y) are indicated (CX-Y). The electron density map is shown next to the model alongside characteristic 2D class averages. (**A**) Negative stain characterization of HALs. Ring diameters are 92 Å, 110 Å, 75 Å, 80 Å, 100 Å, 107 Å, for HALC6_220, HALC24-6_316, HALC20-5_308, HALC25-5_341, HALC18-6_278 and HALC42-7_351, respectively. (**B**) CryoEM characterisation of three large HALs. The ring diameters are 87 Å, 99 Å, and 100 Å for HALC15-5_262, HALC18-6_265, and HALC33-3_343, respectively. Top row left panels: design model colored by chain; Top row, right panels: superpositions of the CryoEM model (gray) and design model (blue). Bottom row: 4.38 Å, 6.51 Å, and 6.32 Å cryoEM electron density maps. Scale bars = 10 nm.

The hallucinated rings are giant structures quite unlike anything in the PDB. The three rings solved by cryoEM, HALC5-15_262, HALC6-18_265 and HALC3-33_343, are 87 Å, 99 Å and 100 Å in diameter and 40 to 50 Å high, with a continuous parallel β-sheet in the lumen of the pore, and outer helices that enforce the curvature and closure of the ring. HALC3-33_343 has a simple helix-loop-sheet structural motif as the repeating unit, while in HALC5-15_262 and HALC6-18_265, the repeating unit contains two distinct helix-loop-sheet elements, which produces an alternating helical outer pattern clearly observable in the 2D class averages. While both structures have reasonable matches to LRRs for their protomers (TM-score of 0.65 for both, but to different structures), the oligomers are strikingly different from any natural protein, with TM-scores of 0.48 and 0.49 respectively (Fig. 4H-I). HALC3-33_343 has an unusual internal loop region breaking the outer helices midway in the repeat, producing a widening of the ring on one side, which is clearly visible in the cryoEM reconstruction; the protomer has a low TM-score (0.48) despite having an LRR-like topology, and the oligomer is even further from anything currently known (TM-score: 0.41) To our knowledge, these designs are the largest cyclic homo-oligomers designed *de novo* to date, and the sophistication of the fold, topology, and high sequence and structural symmetry rivals that in nature: the highest cyclic symmetry recorded in the PDB for naturally occurring proteins is C39 (Vault proteins (*32*), PDB 4HL8 and 7PKY), and there are no closed symmetric α/β ring-like structures.

**Fig. 4.**
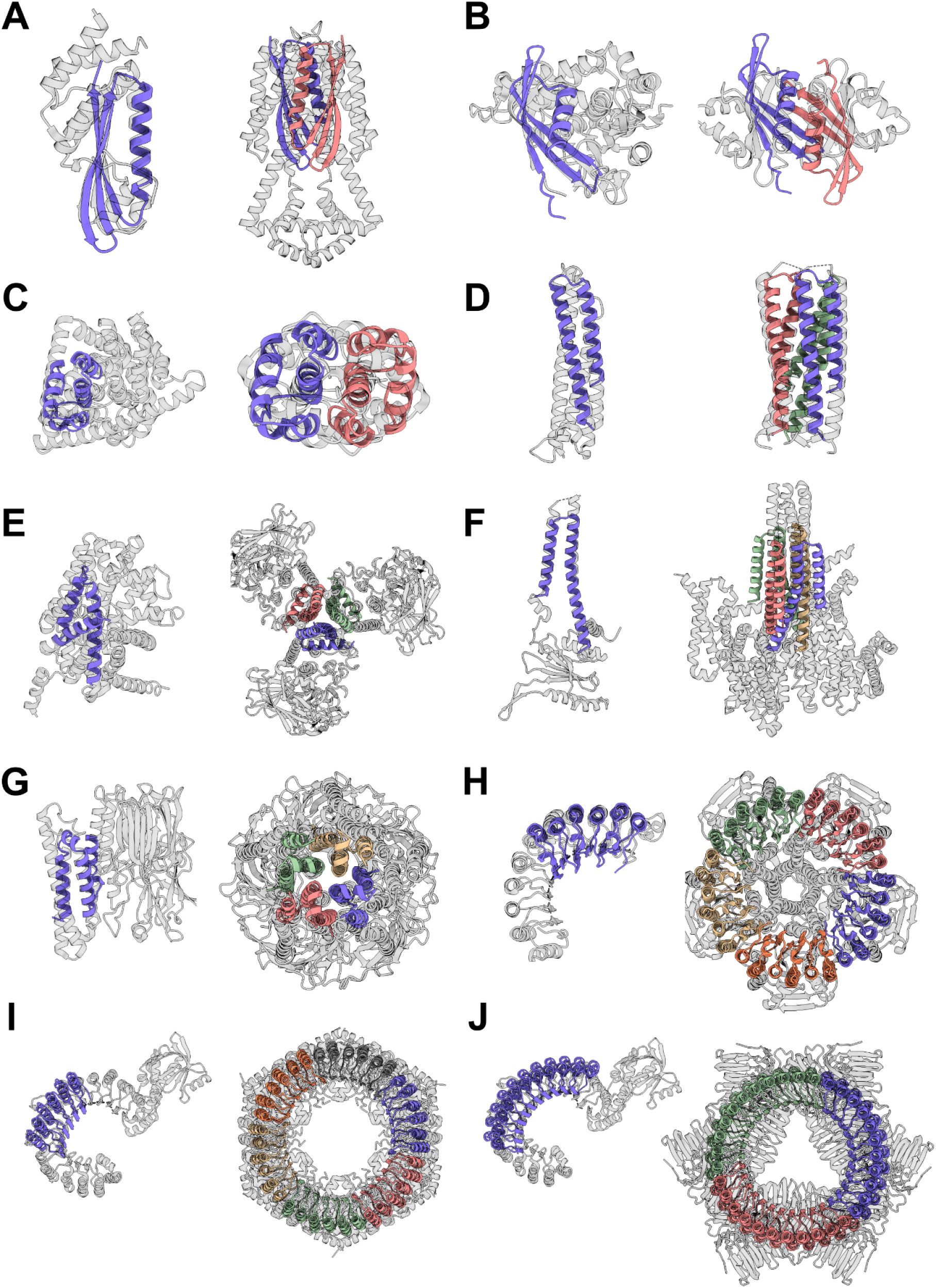
Hallucinated structures differ significantly from their closest matches in the PDB. For each structure solved by crystallography (Fig. 2) or cryoEM (Fig. 3B), the closest structural match to the protomer and to the oligomer are shown on the left and right respectively. Designs are colored by chain and the closest matching PDB is shown in gray. In most cases the closest oligomer has an entirely different structure; this is particularly evident for the larger designs in **G**-**H**. TM-scores (protomer | oligomer) are indicated in parentheses, and the PDB IDs are reported in Table S1. (**A**) HALC2_062 (0.69 | 0.59). (**B**) HALC2_065 (0.67 | 0.54). (**C**) HALC2_068 (0.67 | 0.57). (**D**) HALC3_104 (0.87 | 0.88). (**E**) HALC3_109 (0.78 | 0.69). (**F**) HALC4_135 (0.80 | 0.59). (**G**) HALC4_136 (0.80 | 0.71). (**H**) HALC15-5_262 (0.65 | 0.46). (**I**) HALC18-6_265 (0.65 | 0.49). (**J**) HALC33-3_343 (0.49 | 0.41).

## Conclusion

Our deep learning-based approach to designing cyclic homo-oligomers jointly generates protomers and their oligomeric assemblies without the need for a hierarchical docking approach. We report a rich assortment of *de novo* protein homo-oligomers across the nanoscopic scale, with broad topological diversity while maintaining design constraints such as symmetry and oligomeric state. These hallucinated oligomers differ substantially from natural oligomers in both sequence (median lowest BLAST E-value against UniRef100 of 1.3 for the repeated sequence motifs, Fig. 1D)) and structure (median best TM-scores against the PDB for the protomers and oligomers of 0.67 and 0.57 respectively, Fig. 1C); our computational pipeline interpolates and extends native fold-space rather than simply recapitulating memorized protein structures, demonstrating the power of deep learning to explore previously uncharted regions of the design landscape (Fig. 1B). Our results also highlight the power of the ProteinMPNN method for protein sequence design: of the 30 out of the 192 designs evaluated experimentally by either SEC-MALS, nsEM, cryoEM, or X-ray crystallography, 27 had the intended oligomeric state, and 7 of 19 for which crystallization was attempted formed diffracting crystals (this is a considerably higher crystallization success rate than typical for Rosetta *de novo* designs, and suggests that ProteinMPNN may generate protein surfaces more likely to form crystal contacts).

The high level of abstraction associated with the specification of a loss function enables the design of complex structures with minimal user input, facilitating the design process and making it accessible to non-experts, while generating a rich array of solutions with high experimental success rates. The formalism described here can be extended to other types of complex design tasks, including the design of higher order point group symmetries, arbitrary symmetric or asymmetric hetero-oligomeric assemblies, oligomeric scaffolding of existing functional domains, and design of multiple states, provided a loss function describing the solution can be formalized and computed. Computational requirements and hardware memory limitations become bottlenecks for hallucination of increasingly large structures; the development of computationally less expensive structure prediction methods with fewer parameters, for instance limited to backbone generation, as well as faster-converging algorithms for navigating the sequence space, will further increase the power of the method. Overall, the approach paves the way for the design of larger *de novo* protein structures approaching the complexity found in natural systems, as well as specialty functional nanomaterials integrating catalysis or large multicomponent protein nanomachines(*33*) performing work at the nanoscale.

## Acknowledgements

We thank Ivan Anishchenko, Sergey Ovchinnikov, William Sheffler, Christoffer Norn, Dmitri Zorine, Luki Goldschmidt, and Timothy Huddy for helpful discussions.

## Author contributions

Conceptualization: AC, BIMW, LFM, DB

Methodology: AC, BIMW, LFM, DB

Software: AC, BIMW, LFM, JD, MB, FD, RK

Validation: AC, BIMW, LFM, ST, EK, RJR, AKB

Formal analysis: AC, BIMW, LFM, DB, RJR

Investigation: AC, BIMW, LFM, ST, EK, XL, LC, AKB, AK, HN

Resources: AC, BIMW, LFM, MB, FD, DB

Data curation: AC, BIMW, LFM, DB, AKB, RJR, XL, LC

Writing – original draft: AC, BIMW, LFM, DB

Writing – review & editing: AC, BIMW, LFM, DB

Visualization: AC, BIMW, LFM, RJR

Supervision: DB

Project administration: AC, BIMW, LFM, DB

Funding acquisition: AC, BIMW, LFM, DB

## Data and materials availability

All data is available in the main text or as supplementary materials. Scripts and computational methods are available on GitHub, Crystallographic datasets have been deposited in the PDB (accession codes: 8D03, 8D04, 8D05, 8D06, 8D07, 8D08 8D09). EM maps have been deposited in the EMDB (accession codes: EMD-XXXXX, EMD-XXXXX, EMD-XXXX).

## Supplementary Materials for

### Other Supplementary Materials for this manuscript include the following

Dataframe containing all protein information
Oligomer hallucination code
Design models

## Materials and Methods

### Computational design strategy

We reasoned that the ability of AF2 to predict oligomers could be employed to design such structures using a MCMC search in sequence space in combination with a suitable loss function. The advantage of such a method is its ability to jointly optimize the protomer and oligomer structures, without putting any constraints on the nature of the protomer itself (e.g. the requirement to adopt a well-folded structure in isolation as is typically the case for docking approaches). We employed simplifications during AF2 predictions to reduce computational cost, and defined a composite loss function composed of structure quality terms and a geometric term.

MCMC trajectories were initialized with a random protomer sequence of specified length, with the composition of amino acids respecting the BLOSUM62 background frequencies. Cysteines were disallowed for all hallucinations. Protomers sequences were concatenated to generate oligomeric assemblies during AF2 prediction: chain breaks in the concatenated protomer sequences were specified by re-indexing residues after the break with a 200 increment, resulting in AF2 predicting them as separate chains. To reduce computational costs the number of recycles was set to 1, the number of ensembles was also set to 1, and AMBER relax was not performed. After each prediction losses were computed on the AF2 prediction confidence metrics (pLDDT, pTM, pAE) as well as the coordinates of the predicted structure.

Mean AF2 pLDDT and AF2 pTM scale between 0 and 1, where higher values are better, thus the loss (by definition the objective to minimize) was calculated for each as one minus their respective values. For enforcing cyclic symmetry we computed a cyclic loss term defined as the standard deviation between the center of mass of adjacent protomers (computed on Cα). Minimizing this value enforces cyclic symmetry.

The loss functions computed to generate all cyclic oligomers <= C7 was:

`Dual_cyclic: loss = 1 - 0.5*(AF2pTM +AF2 pLDDT) + standard deviation(center of masses)`

After an initial prediction, mutations were introduced in the protomer sequences (tied positions), and the structure re-predicted. Positions with low pLDDT values (lowest half) were targeted, and mutations were chosen based on the BLOSUM62 substitution frequencies. The number of mutations at each step was linearly decayed over the course of the trajectory starting from 3 per protomer down to 1.

Simulated annealing was employed during optimization, with the starting temperature set to 0.01 and the half-life of the exponential decay set to 500 steps. Mutations were accepted or rejected according to the Metropolis criterion

All parameters and loss functions mentioned above and several others are available in the code repository. We highlight that the command to generate e.g. a homo-trimer is simply:

`./oligomer_hallucination.py --L 65 --oligo AAA+ --loss dual_cyclic --out C3_oligomer`

Modest computational means were sufficient to hallucinate assemblies up to C7 with protomer lengths of 65 amino acids. The largest C7 assemblies required a week on a single CPU with 6 GB of memory to generate 300 steps, which can be sufficient for convergence (pLDDT > 0.70 and pTM > 0.70) . For smaller assemblies (e.g. a C3 with protomers composed of 65 amino acids) approximately 500 steps per day could be obtained on a single CPU with 5 GB of memory.

The structures generated from AF2 hallucination were sequence re-designed with ProteinMPNN using only the restrictions that protomer sequences in the oligomeric assembly were tied to be identical, and cysteines were disallowed. For each backbone 24-48 sequences were generated with ProteinMPNN using a temperature of 0.2. The quality of these sequences was assessed with AF2 using all 5 models (model_1-5_ptm), checking both the confidence metrics and the structural recapitulation of the original backbone geometry. Sequences were filtered on having AF2 pLDDT > 0.75, and a RMSD to the original protomer backbone < 1.5 Å (computed with TMalign, (*34, 35*)). For each original backbone the four designs with highest AF2 pLDDT were inspected by eye, and up to three MPNN sequences per original input backbone were ordered for experimental testing.

All code is available on GitHub: to be uploaded
ProteinMPNN is available on GitHub: to be uploaded
AF2 is available at: https://github.com/deepmind/alphafold
RoseTTAFold is available at: https://github.com/RosettaCommons/RoseTTAFold

### RoseTTAFold prediction of oligomers

An updated version of RoseTTAFold was used to evaluate designed oligomers. This RoseTTAFold model has multiple architectural improvements over the original published model, including; 1) use of a 3D track from the beginning, with coordinates from a template or the previous recycling round, 2) communication between 1D, 2D, and 3D tracks through attention biasing, and 3) use of recycling that executes the network multiple times with the updated input embeddings based on outputs from the previous cycle. The model was trained with 3 recycling steps. The training dataset comprised; 1) both single-chain and biologically relevant complex structures from the PDB released before April 30, 2020, and 2) AlphaFold2 model structures for UniRef50 representatives. For the examples used during training that were oligomers, we added 200 to the residue numbers of the following subunits to indicate chain breaks to the network. Two rounds of model training were performed; 1) an initial training (200 epochs, with 25600 examples per epoch and a batch size of 64) based on the masked language recovery loss, distogram prediction loss, predicted LDDT loss, and FAPE loss followed by, 2) fine-tuning (50 epochs, with 25600 examples per epoch and a batch size of 64) with additional loss terms on bond geometry and van der Waals scoring function. We trained the model with a crop size of 256 residues, and then fine-tuned it with a larger crop (384 residues). The AdamW Optimizer with default pytorch parameters was used. For the initial training we linearly increased the learning rate to 0.001 over the first 1000 optimization steps, and further decreased the learning rate by a factor of 0.95 for every additional 5000 optimization steps. The fine-tuning stage started from the pre-trained model weights, and used the lower learning rate (0.0005), no warm-up steps, and the same step-wise learning rate decay.

During inference we added 200 to the residue indices of subsequent subunits to indicate chain breaks, as we did during model training. The model was recycled 20 times, and the predicted structure having the highest LDDT estimation was selected. The oligomer structure predictions were generated from the designed sequence only, without any MSA or template information.

### Comparison to natural proteins

The outputs generated during AF2 hallucination and ProteinMPNN re-design were assessed for their sequence and structure novelty. Sequence homologues were searched using BLAST (Protein-Protein BLAST version 2.11.0+) against UniRef100 (snapshot from March 2, 2022) and the E-value of the best hit reported. Both the sequence of the protomer as well as the repeated sequence motif were queried. In the case of small HALs, the protomer and repeated sequence motif were equivalent, but not in the case of large HALs (i.e. HALCX-Y), where protomers are composed of multiple repeated sequence motifs. Structural comparisons to published structures were performed at the protomer level (using TMalign version 20190425) against the PDB (snapshot from April 15, 2022) and over the whole oligomer (using MMalign version 20210816) against all biounits assigned in the PDB (snapshot from April 15, 2022). In both cases results are reported as TM-score.

### Representation of the structural space

A representation of the structural space covered by the outputs of the hallucination trajectories compared to all *de novo* cyclic structures deposited in the PDB is shown in Fig. 1B. The plot was obtained by Multidimensional scaling (as implemented in the sklearn python library) on a pre-computed pairwise distance matrix. Pairwise distances were defined as 1-TM-score, and the score computed with TMalign (version 20190425). The list of 162 *de novo* cyclic structures was obtained by using the following gate on a snapshot of the PDB from April 17, 2022:

`Entry Polymer Composition == homomeric protein &`
`Polymer Entity Sequence Length >= 40 &`
`Structure Keywords contains ’de novo’ &`
`Type == Cyclic`

1ec5,1g6u,1jm0,1jmb,1lt1,1mft,1ovr,1ovu,1ovv,1u7j,1u7m,1uw1,1vjq,1y47,1y66,2gjf,2gjh,2i7u,2jst,2kik,2mg4,2p05,2p09,2wqh,2zgd,2zgg,3cwo,3dgo,3lt8,3lt9,3lta,3ltb,3ltc,3ltd,3m22,3m24,3mlg,3ol0,3rhu,3tdm,3tdn,3v1b,3v1c,3v1d,3v1e,3v1f,3vjf,3ww7,3ww8,3wwb,3wwf,4db8,4dba,4etj,4f2v,4glu,4hxt,4loa,4lpu,4lpv,4lpw,4lpx,4lpy,4m6a,4ndj,4ndk,4ney,4nez,4o60,4ow4,4pww,4qfv,4rjv,4wpy,4yfo,4yxy,4zcn,4zxz,5a0o,5bvb,5c39,5di5,5dn0,5dns,5dqa,5dra,5dzb,5eil,5f53,5h78,5hpn,5hry,5hrz,5hs0,5i1z,5izs,5j0h,5j0i,5j0j,5j0k,5j0l,5j10,5j2l,5j73,5k7v,5kay,5kba,5kwd,5l0p,5od9,5tph,5u35,5vl4,5ys7,6ff6,6g6q,6idc,6iei,6kos,6m6z,6msq,6msr,6n9h,6naf,6nek,6nla,6nx2,6ny8,6nye,6nyi,6nyk,6nz1,6nz3,6o0c,6o0i,6o35,6qsh,6tjb,6tjc,6tjd,6u1s,6v8e,6veh,6w40,6w6x,6wxo,6wxp,6xh5,6xi6,6xns,6xr2,6xss,6xt4,6y7n,6zv9,7ax0,7bww,7dns,7k3h,7kxs,7m0q,7nbi

### Plasmid construction

Plasmids for expressing HALs were constructed from synthetic DNA according to the following procedure: Linear DNA fragments (Integrated DNA Technologies, IDT eblocks) encoding design sequences and including overhangs suitable for a BsaI restriction digest were cloned into custom target vectors using Golden Gate Assembly. All subcloning reactions resulted in C-terminally HIS-tagged constructs, either as: MSG-design-GSHHHHHH (entry vector LM670) or a MSG-design-GS<u>GSHHW</u>GSTHHHHHH (entry vector LM627), where the underlined sequence is the SNAC-tag used for cleaving the HIS-tag for crystallization (*36*).

The entry vectors for Golden Gate cloning are modified pET29b+ vectors that contain a lethal ccdb gene between the BsaI restriction sites that is both under control of a constitutive promoter and in the T7 reading frame. The lethal gene reduces background by ensuring that plasmids that do not contain an insert (and therefore still carry the lethal gene) kill transformants. The vectors were propagated in ccdb resistant NEB Stable cells (New England biolabs C3040H, always grown from fresh transformants). Plasmids were deposited with Addgene.

Golden Gate reactions (5 uL per well) were set up on a 96 well PCR plate as:

10x T4 Buffer 0.5 uL 10x T4 Buffer (New England Biolabs B0202S)
Vector 10-20 fmol Vector (either LM627 or LM670)
BsaI-HFv2 3U 0.15 uL BsaI-HFv2 (New England Biolabs R3733L)
T4 Ligase 100U 0.25uL T4 Ligase (New England Biolabs M0202L)
+ (20-40 fmol) linear DNA fragment, typically 1 uL of 10 ng/uL stock
Complete with nuclease-free water to 5 uL total reaction volume.

The reactions were incubated at 37 °C for 20 minutes, followed by 5 min at 60 °C in a thermocycler (Biorad T100) with the lid heated to 105 °C.

### Small-scale protein solubility screen

For initial solubility screens, Golden Gate reaction mixtures were transformed into BL21(DE3) (New England Biolabs) as follows: 1 uL of reaction mixture was added to 6-8 uL of competent cells on ice in a 96 well PCR plate. The mixture was incubated on ice for 30 minutes, then heat-shocked for 10 s at 42 °C in a block heater (IKA Dry Block Heater 3), then rested on ice for 2 minutes. Subsequently, 100 uL of room temperature SOC media (New England Biolabs) was added to the cells, followed by incubation at 37 °C with shaking at 1000 rpm on a Heidolph Titramax1000 / Incubator 1000.

The transformations were then grown in a 96 well deep-well plate (2 mL total well volume) in autoclaved LB media supplemented with 50 μg mL^-1^ Kanamycin at 37 °C and 1000 rpm. In the following protocols all growth plates were covered with breathable film (Breathe Easier, Diversified Biotech) during incubation.

The following day, glycerol stocks were made from the overnight cultures (100 uL of 50 % [v/v] Glycerol in water mixed with 100 uL bacterial culture, frozen and kept at -80 °C. Subsequently, two 96 deep well plates were prepared with 900 uL per well of autoclaved Terrific Broth II (MP biomedicals) supplemented with 50 μg mL^-1^ Kanamycin, and 100 uL of the overnight culture were added and grown for 1.5 h at 37 °C, 1200 rpm (Heidolph Titramax1000 / Incubator 1000). The cultures were then induced with IPTG by adding 10 uL of 100 mM (final concentration approximately 1 mM) per well with an electric repeater pipette (Eppendorf, E4x series), and grown for another 4 h at 37 °C, 1200 rpm. Cultures were combined into a single 96 well plate for a total culture volume of 2 mL and harvested by centrifugation at 4000 x g for 5 min. Growth media was discarded by rapidly inverting the plate, and harvested cell pellets were either processed directly, or frozen at -80 °C.

Proteins were purified by HIS tag-based Immobilized metal affinity chromatography (IMAC). Bacterial pellets were resuspended and lysed in 300 uL B-PER chemical lysis buffer (Thermo Fisher Scientific) supplemented with 0.1 mg mL^-1^ Lysozyme (from a 100 mg mL^-1^ stock in 50 % [v/v] Glycerol, kept at -20 °C, Millipore Sigma), 50 Units of Benzonase per mL (Merck/Millipore Sigma, stored at - 20 °C), and 1 mM PMSF (Roche Diagnostics, from a 100 mM stock kept in Propan-2-ol, stored at room temperature). The plate was sealed with an aluminum foil cover and vortexed for several minutes until the bacterial pellet was completely resuspended (on a Vortex Genie II, Scientific Industries). The lysate was incubated, shaking for 5 minutes, before being spun down at 4000 x g for 15 minutes. In the meantime, 75 uL of Nickel-NTA resin bed volume (Thermo Scientific, resin was regenerated before each run and stored in 20 % [v/v] Ethanol) was added to each well of a 96 well fritted plate (25 μm frit, Agilent 200953-100). To increase wash step speed, the resin was equilibrated on a plate vacuum manifold (Supelco, Sigma) by drawing 3 x 400 uL of Wash buffer (20 mM Tris, 300 mM NaCl, 25 mM Imidazole, pH 8.0) over the resin using the vacuum manifold at its lowest pressure setting.

The supernatant (280 uL) of the lysate was extracted after the spin down and applied to the equilibrated resin and allowed to slowly drip through over ∼5 minutes. Subsequently the resin was washed on the vacuum manifold with 3 x 400 uL of Wash buffer. Lastly the fritted plate spouts were blotted on paper towels to drain excess Wash buffer. Then 250 uL of Elution buffer (20 mM Tris, 300 mM NaCl, 500 mM Imidazole, pH 8.0) was applied to each well and incubated for 5 minutes before eluting the protein by centrifugation at 1500 x g for 5 minutes into a 96 well collection plate. Eluate was stored at 4 °C.

Screening samples for EM and initial SDS-PAGE (Biorad Criterion 26well stain free - anykD) analysis to assess solubility were prepared using this method. Correct protomer masses were verified by Liquid chromatography-mass spectrometry (LC-MS, Agilent) on soluble eluates. To identify the molecular mass of each protein, intact mass spectra was obtained via reverse-phase LC/MS with an Agilent G6230B TOF on an AdvanceBio RP-Desalting column (A: H2O with 0.1% Formic Acid, B: Acetonitrile with 0.1% Formic Acid), and subsequently deconvoluted with Bioconfirm using a total entropy algorithm.

### Larger-scale protein expression and purification for biophysical studies

Overnight autoinduction cultures were seeded from the glycerol stocks made for the small scale screen. Growth media was TB-II autoinduction media: TB-II (Terrific Broth II, MP biomedicals - prepared according to manufacturer’s specifications: 50 g / L, autoclaved) supplemented with Studier 5052 components from a 50x stock (final concentrations: 5 g / L glycerol, 0.5 g / L dextrose, 2 g / L lactose monohydrate), and 2 mM MgSO_4_.

For the initial screen of 150 AF2 hallucinations, 50 mL cultures were grown in 250 mL baffled flasks (24h, 37 °C, 250 rpm). For the subsequent screen of the MPNN designed sequences, 15 mL cultures were grown in 125 mL baffled flasks (16h, 37 °C, 250 rpm). Cultures were harvested by centrifugation at 4000 x g for 5 minutes, and pellets were stored frozen at -80 °C, or processed directly.

The parameters for the purification of the initial 150 AF2 based hallucinations and the MPNN redesigned sequences are given as ( AF2 | MPNN ) differed slightly because of differences in expression culture volume ( 50 mL | 15 mL )

For protein purification, pellets were resuspended in ( 10 mL | 5 mL ) Wash buffer (20 mM Tris, 300 mM NaCl, 25 mM Imidazole, pH 8.0 at room temperature, supplemented with 0.1 mg mL^-1^ Lysozyme, 0.01 mg mL^-1^, Deoxyribonuclease I (DNAse I, Millipore Sigma), 1 mM PMSF) by vortexing for several minutes until the pellet was fully resuspended. The resuspension was sonicated (Qsonica, Q500 with a: 4 pronged horn | 24 pronged horn) as 10 s ON, 10 s OFF, (45% | 80 %) amplitude for 5 minutes of total ON time, and samples were kept on ice during the whole procedure.

The sonicated lysate was centrifuged at (14000 x g | 4000 x g) for 15-45 minutes to remove the insoluble fraction. Plates with 25 μm bottom frits with ( 24 | 48 ) wells (Agilent 201415-100 | 201003-100) were filled with ( 1 mL | 0.5 mL ) of bed Ni-NTA resin (Qiagen or Thermo Fisher), and equilibrated with three rinses of Wash buffer (at least 30 resin bed volumes) on a vacuum manifold as described above.

The fritted plate spouts were closed with parafilm, and the supernatant was added to each well. The plate was sealed and incubated lightly agitated for 30 minutes. The supernatant was drained from the resin, and the resin bed washed three times with ( 10 mL | 5 mL ) of Wash buffer (at least 30 resin bed volumes) on the vacuum manifold. Excess Wash buffer was blotted from the spouts on paper towels, and the resin was pre-eluted with 80% resin bed volume of Elution buffer, followed by protein elution into ( 1.1 mL | 0.8 mL ) of Elution buffer (20 mM Tris, 300 mM NaCl, 500 mM Imidazole, pH 8.0).

### Size Exclusion Chromatography (SEC)

IMAC eluates were sterile-filtered through a 96 well filter plate (0.2 μm polyethersulphone (PES) membrane, Agilent 204510-100) by centrifugation at 2000 x g for 5 minutes.

Size exclusion chromatography was performed using an autosampler-equipped Akta pure system (Cytiva) on a Superdex S200 Increase 10/300 GL column at room temperature. The running buffer was 20 mM Na-PO4, 100 mM NaCl, pH 7.4 at room temperature. Selected fractions (shown in Figure S4) were pooled and concentrated using Spin filters (3 kDa molecular weight cutoff, Amicon, Millipore Sigma) and stored at 4 °C before downstream characterizations. Protein identities were confirmed by reverse-phase LC-MS as described above.

SEC retention volume to molecular weight equivalencies were calibrated with protein standards (Cytiva LMW and HMW kits for the S75 and S200 columns, respectively).

Samples for electron Microscopy were purified by SEC using a Superdex 6 10/300 GL increase column (Cytiva) and TBS running buffer (25 mM Tris pH 8.0, 100 mM NaCl). SEC elution fractions corresponding to the design’s theoretical elution volumes were concentrated in TBS prior to structural and biochemical analysis.

### Size Exclusion Chromatography - Multi Angle Light Scattering (SEC-MALS)

Pooled SEC samples were analyzed by SEC-MALS in 20 mM Na-PO4, 100 mM NaCl, pH 7.4 on a Superdex 75 10/300 or Superdex 200 10/300 column in line with a Heleos multi-angle static light scattering and an Optilab T-rEX detector (Wyatt Technology Corporation). Data was analyzed using ASTRA (Wyatt Technologies) to calculate the weighted average molar mass (Mw) of the selected species and the number average molar mass (Mn) to determine monodispersity by polydispersity index (PDI) = Mw/Mn.

### Circular Dichroism (CD)

Circular Dichroism was performed on a Jasco 1500 CD spectrometer with a 6 sample rotating turret. Samples were placed in 1 mm pathlength cuvettes (Hellma QS Quartz cell) at concentrations of 0.25 mg mL^-1^ in 20 mM Na-PO4, 100 mM NaCl, pH 7.4 buffer. The temperature was ramped from 25 °C to 95 °C, recording full CD spectra between 200 and 260 nm in 10 °C intervals, and reading at 222 nm in 2 °C intervals. After reaching 95 °C the samples were allowed to cool back to 25 °C before recording a final spectrum. Samples were recovered, filtered over a 0.2 μm PES membrane, and re-run over SEC as described above.

### Crystallography sample preparation and data collection

19 designs were chosen to undergo crystallization screens. Each design was expressed as described above in 0.5 L cultures. Following affinity purification, each design underwent SEC into SNAC cleavage buffer (100 mM CHES, 100 mM NaCl, 100 mM acetone oxime, 500 mM guanidine HCl, pH 8.6). Following SEC, 2 mM of NiCl_2_ was added and the solution was incubated overnight at 37°C. Following cleavage, the solutions containing the cleaved protein products were incubated with 1 mL Ni-NTA resin to bind any uncleaved product, and the flow through was collected. Following SEC into Crystallization buffer (20mM Tris, 50 mM NaCl, pH 8.0), each sample was concentrated to approximately 15 mg mL^-1^. The following sitting drop broad screens were set up at room temperature with three protein:crystallization condition ratios (1:1, 1:2, 2:1) using the mosquito pipetting instrument (sptlabtech): Midas (Molecular Dimensions), Proplex (Molecular Dimensions), JCSG+ (Molecular Dimensions), Morpheus (Molecular Dimensions), Pact Premier (Molecular Dimensions), LMB (Molecular Dimensions), Index (Hampton Research) and PGA (Molecular Dimensions). Each was monitored weekly for crystal growth using the JANSi UVEX imaging system.

The following conditions yielded diffracting crystals for our designs: 0.05 M Cesium chloride, 0.1 M MES pH 6.5, 30% Jeffamine M-600 (HALC3_104); Morpheus condition H5 (HALC3_109); 0.1 M BIS-TRIS pH 6.5, 2.0 M Ammonium sulfate (HALC2_062); 0.2 M Lithium sulfate monohydrate, 0.1 M BIS-TRIS pH 6.5, 25% w/v Polyethylene glycol 3,350 (HALC4_135); 0.1M SPG buffer pH 5 25% w/v PEG 1500 (HALC4_136), 0.04 M Potassium phosphate, 16% PEG 8000, 20% Glycerol (HALC2_068); and 0.2 M Ammonium nitrate pH 6.3, 20% PEG 3350 (HALC2_065). Where required, crystals were cryoprotected with 20% glycerol or 25% ethylene glycol prior to flash freezing in liquid nitrogen. Data collection was done using the Advanced Photon Source synchrotron. Images were integrated using XDS 20220110 (*37*). Aimless (*38*) was used for scaling and merging. Phaser 2.8 (*39*) was used for molecular replacement using the design models as search models (either monomer or oligomeric complex). Models were built using Coot 0.9.8 (*40*) and refined with Phenix refine from Phenix 1.20 (*41*) and RefMac (*42*) from CCP4 7.1 (*38*) suite. All structures were validated using MolProbity 4.5.1(*43*). Crystallographic statistics are available in Table S3.

### Negative Stain Electron Microscopy (nsEM)

SEC fractions corresponding to the designs were concentrated in TBS prior to negative stain EM screening. Samples were then immediately diluted 5 to 150 times in TBS buffer (25 mM Tris, 100 mM NaCl, pH 8.0) depending on the concentration of the samples. A final volume of 5 μL was applied on negatively glow discharged, carbon-coated 400-mesh copper grids (01844-F, TedPella Inc.), then washed with Milli-Q Water and stained using 0.75% uranyl formate as previously described (*44*). Air-dried grids were then imaged on either a FEI Talos L120C TEM (FEI Thermo Scientific) equipped with a 4K × 4K Gatan OneView camera at a magnification of 57,000x and pixel size of 2.51 Å. Micrographs collection was automated using EPU software (FEI Thermo Scientific) and were imported into CisTEM software (*45*) or cryoSPARC software (*46, 47*). CTF estimation was done with CTFFIND4 and a circular blob picker was used to select particles which were then subjected to 2D classification. *Ab initio* reconstruction and homogeneous refinement in Cn symmetry were used to generate 3D electron density maps. All EM maps can be found in supplementary data.

### CryoEM Sample Preparation and Data Collection

CryoEM grids were prepared by diluting protein samples with TBS 1 to 10 times immediately before applying 3.5 μL to glow-discharged 400 mesh, C-flat, 2 micron holes, 2 micron spacing, CF-2/2-4C (CF-224C-100) (Electron Microscopy Sciences) cryoEM grids. For some samples, multiple blots were applied in order to obtain the best particle density. All grids were blotted using a blot force of 0 and 5.5 second blot time at 100% humidity and 4°C and plunge-frozen in liquid ethane using a Vitrobot Mark IV (FEI Thermo Scientific). All cryoEM grids were screened on a Glacios transmission electron microscope (FEI Thermo Scientific) operated at 200 kV and equipped with a Gatan K2 or K3 Summit direct detector. Automated glacios data collection was carried out using Leginon (*48*) at a nominal magnification of 36,000x (1.16 Å/pixel). Movies were acquired in counting mode fractionated in 50 frames of 200 ms at 8.5 e-/pixel/sec for a total dose of ∼65e-/Å^2^. Details of data processing for each design are illustrated in Figures S9-S11.

### CryoEM data processing

Multiple datasets were collected for each design and combined early on during processing. See Fig. S9-S11 and processing flowcharts for details. Briefly, images were manually curated to remove poor quality acquisitions such as bad ice or large regions of carbon. Dose-weighting and image alignment of all 50 frames was carried out using MotionCor2 (*49*) with 5X5 patch or with cryosparc v2 patch alignment tool with default parameters. Super-resolution data was binned 2X during alignment. Initial CTF parameters were estimated using CTFfind4 (*50*). Particle picking was done with a gaussian blob picker and in some cases followed by a template picker. Particles were extensively classified in 2D to remove ice and noisy particles, yielding in some cases relatively few particles. Starting models for all designs were always obtained *ab initio*, despite clear evidence of the expected design in 2D. FSC curves were generated using cryoSPARC. All EM maps have been deposited in the EMDB (accession codes: EMD-XXXXX, EMD-XXXXX, EMD-XXXXX), and can be found in the supplementary data.

### Visualization and figures

All structural images for figures were generated using PyMOL, Chimera or ChimeraX. Data was processed and figures were plotted using Pandas, MatplotLib, and Seaborn python libraries. Figures were further rendered and assembled using Adobe Illustrator and Inkscape.

---

**Fig. S1.**
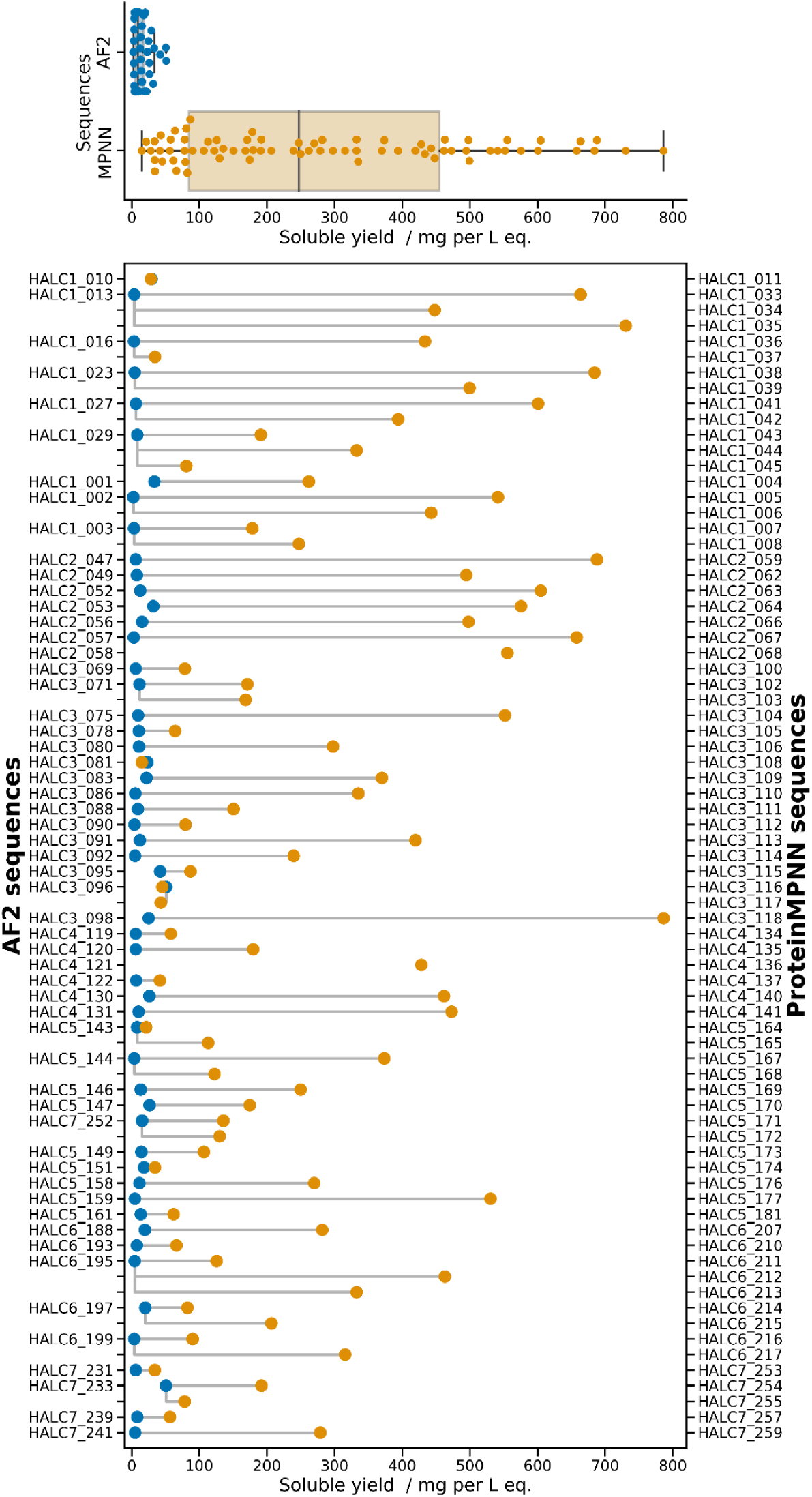
Soluble yield of AF2 and MPNN designed sequences for small HALs. Bottom plot shows the total soluble protein yield per liter equivalent calculated from integrating the SEC traces (and normalizing by the sequence-specific extinction coefficients) for the original AF2 designs, compared to their MPNN redesigns. In some cases more than one MPNN sequence per backbone was ordered. The top plot summarizes the difference in yield: for the AF2 designs a median yield of 9 mg per L eq. as compared to 247 mg per L eq. for the MPNN sequences.

**Fig. S2.**
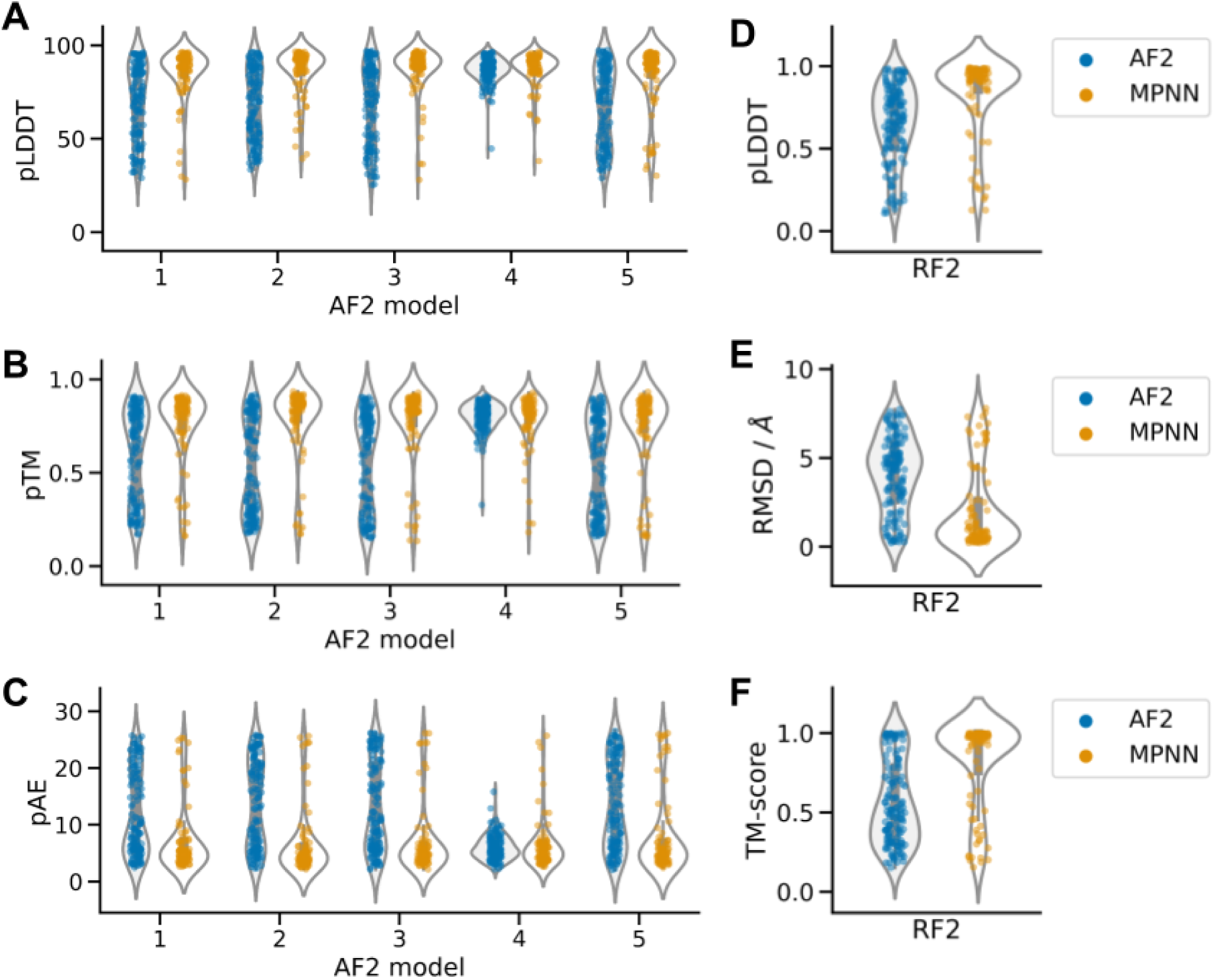
AF2 sequences are not well cross-validated, unlike ProteinMPNN ones. **A**-**C** AF2 predictions of AF2-generated sequences indicate high prediction confidence for the model used during design (model_4_ptm), but low confidence for all other models. In contrast, ProteinMPNN-designed sequences predict better across all five models. **E**-**F** RF2 predictions compared to the designed structure. (**D**) RF2 fails to predict AF2 designed sequences reliably, but confidently predicts ProteinMPNN sequences. **E**-**F** RF2 models of ProteinMPNN sequences are close to their designs, while models predicted from the sequence of AF2 hallucinations fail to recapitulate the designed structure.

**Fig. S3.**
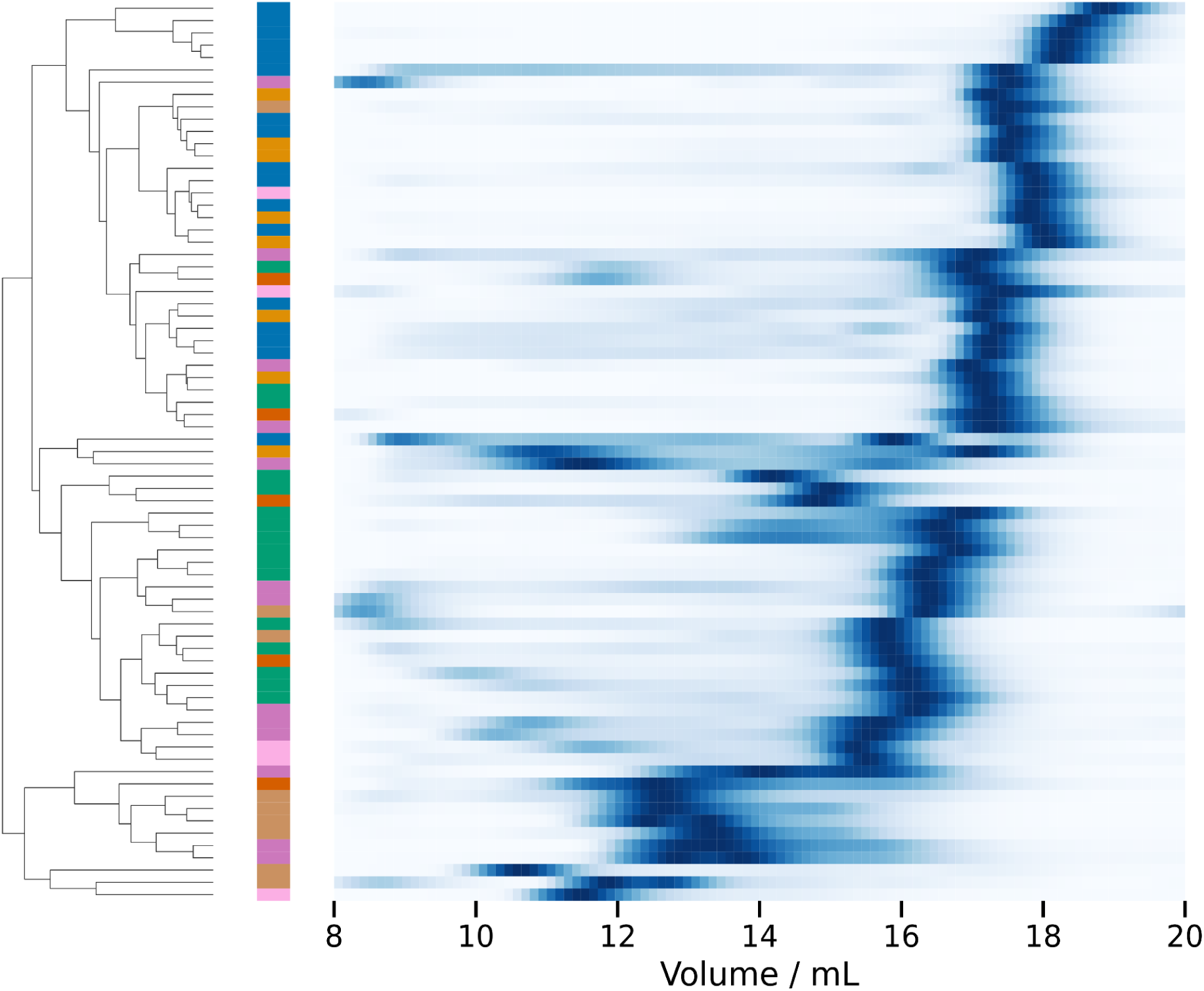
Normalized SEC elution profiles of small HALs. Samples were run on a Superdex 200 increase 10/300 GL following IMAC purification. The results are shown clustered by profile similarities (using Euclidean distances), and for each, the designed oligomeric state is indicated by color (C1; blue, C2; orange, C3; green, C4; red, C5; dark pink, C6; brown, C7; light pink).

**Fig. S4.**
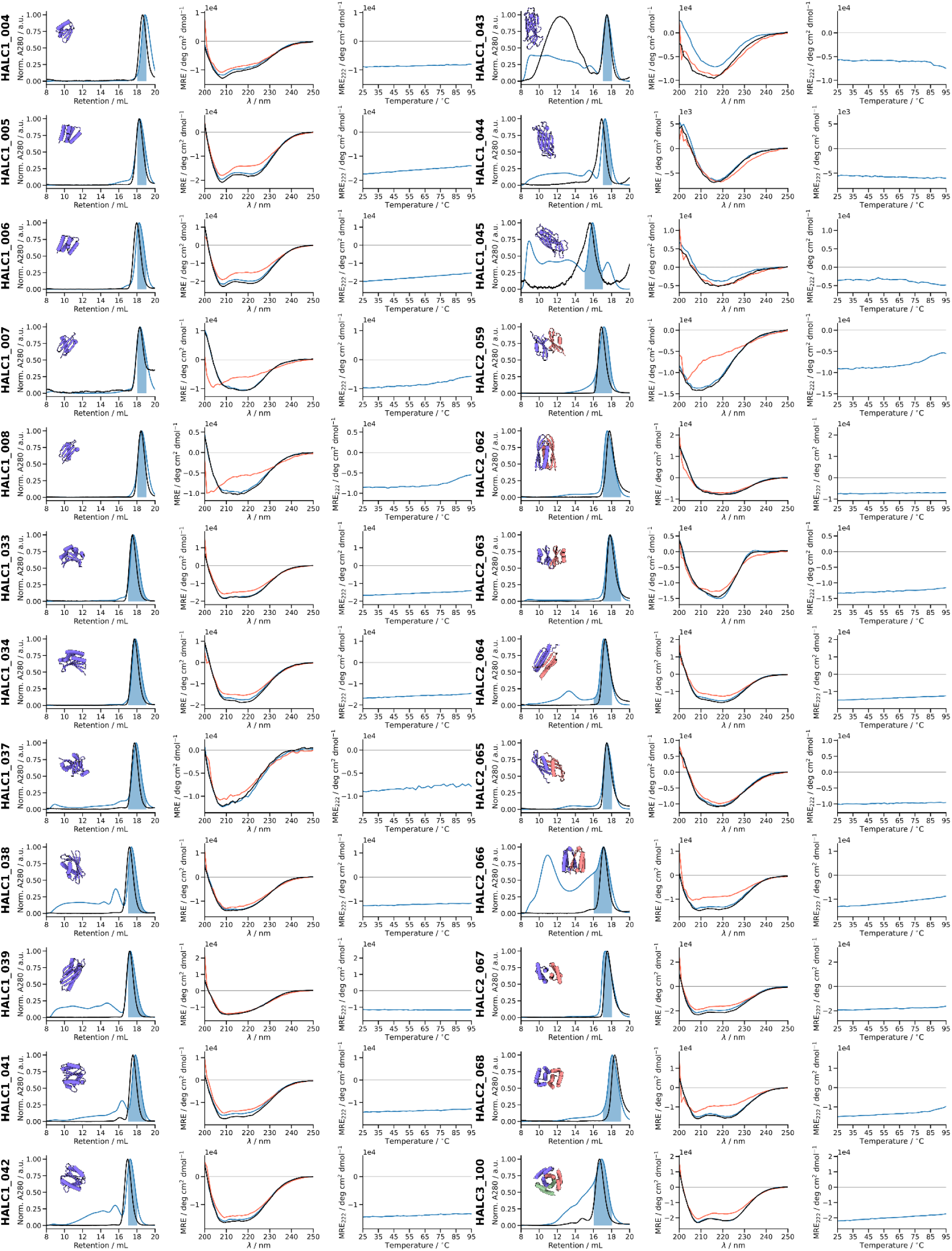
Characterization of HALC1_004 to HALC3_100. The first column shows the SEC elution profile (Superdex 200 increase 10/300 GL) after IMAC (blue, collected fractions indicated by the shaded region), and after heating the sample to 95 °C (black). The second column shows the CD spectra at 25 °C (blue), at 95 °C (red) and after cooling back to 25 °C (black). The third column shows the circular dichroic signal at 222 nm during temperature ramping.

**Fig. S4.**
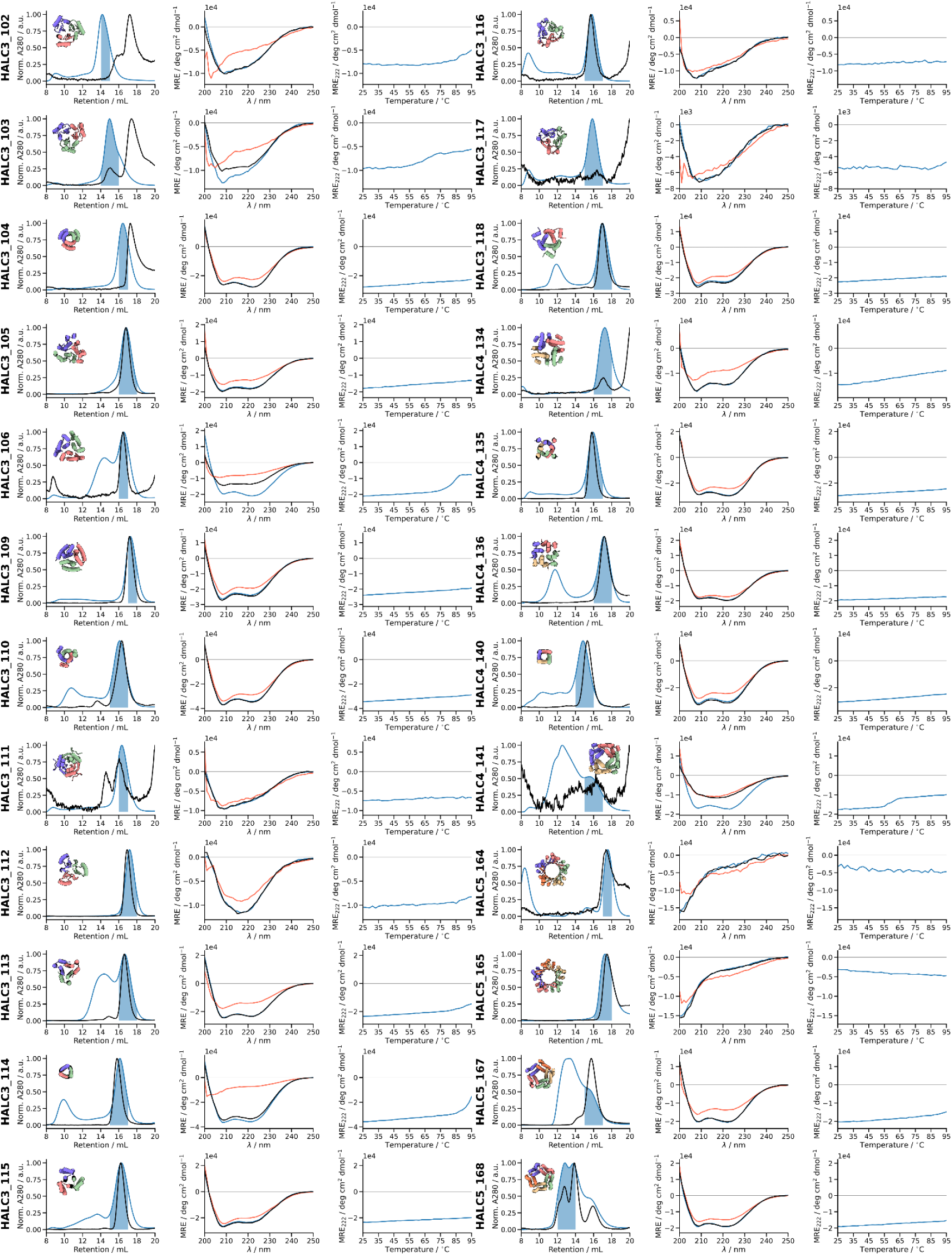
Characterization of HALC3_102 to HALC5_168. The first column shows the SEC elution profile (Superdex 200 increase 10/300 GL) after IMAC (blue, collected fractions indicated by the shaded region), and after heating the sample to 95 °C (black). The second column shows the CD spectra at 25 °C (blue), at 95 °C (red) and after cooling back to 25 °C (black). The third column shows the circular dichroic signal at 222 nm during temperature ramping.

**Fig. S4.**
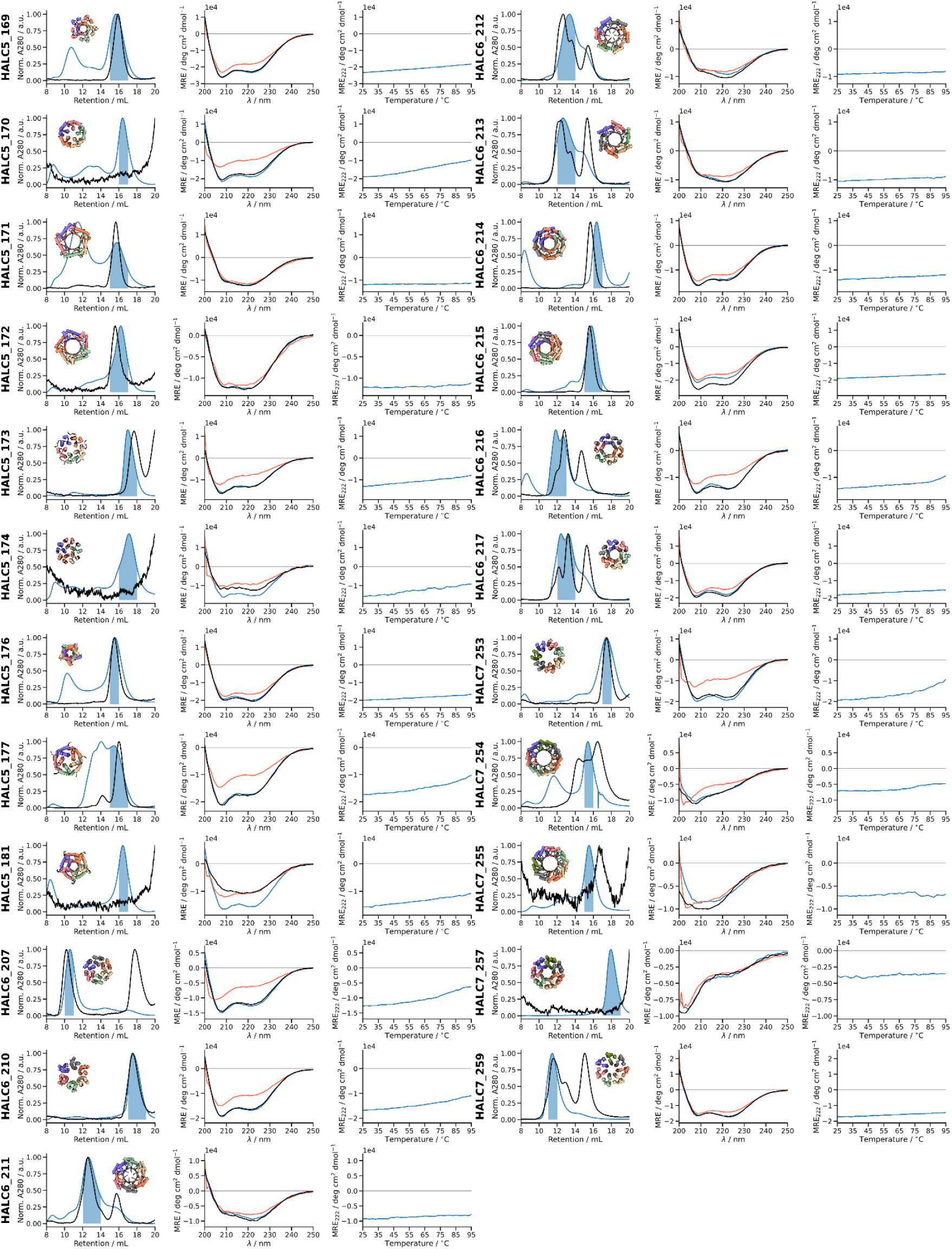
Characterization of HALC5_169 to HALC7_259. The first column shows the SEC elution profile (Superdex 200 increase 10/300 GL) after IMAC (blue, collected fractions indicated by the shaded region), and after heating the sample to 95 °C (black). The second column shows the CD spectra at 25 °C (blue), at 95 °C (red) and after cooling back to 25 °C (black). The third column shows the circular dichroic signal at 222 nm during temperature ramping.

**Fig. S5.**
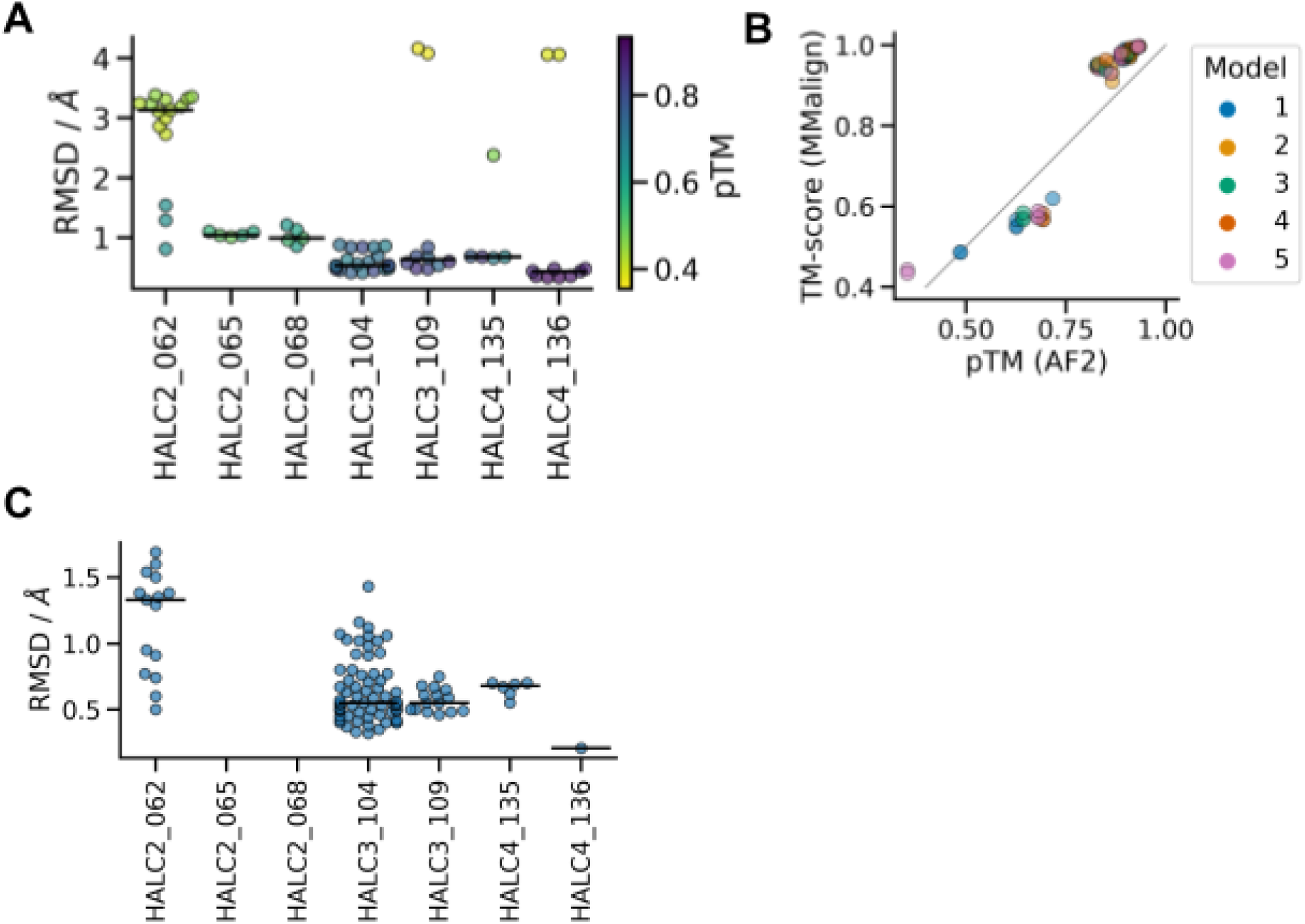
Comparison between AF2 models and crystallographic structures. (**A**) For each design, five models (one for each _ptm model, 10 recycles) were compared to the biounit. If multiple biounits were present, alignments against all bionunits are shown. Alignments were generated using MMalign, and the median RMSD for each design is indicated by a horizontal line. Models that were more confidently predicted (higher pTM values) were closer to the experimentally-validated structures as shown by the color bar. (**B**) The pTM value from each AF2 model correlates with the actual TM-score (from MMalign) between design and structures. The parity is indicated by a grey line. (**C**) Structural matching between chains of the asymmetric unit of each design. Pairwise alignments and RMSD values were computed with TMalign, and the median is indicated by a horizontal line. Designs lacking data points only contained one chain in the asymmetric unit.

**Fig. S6.**
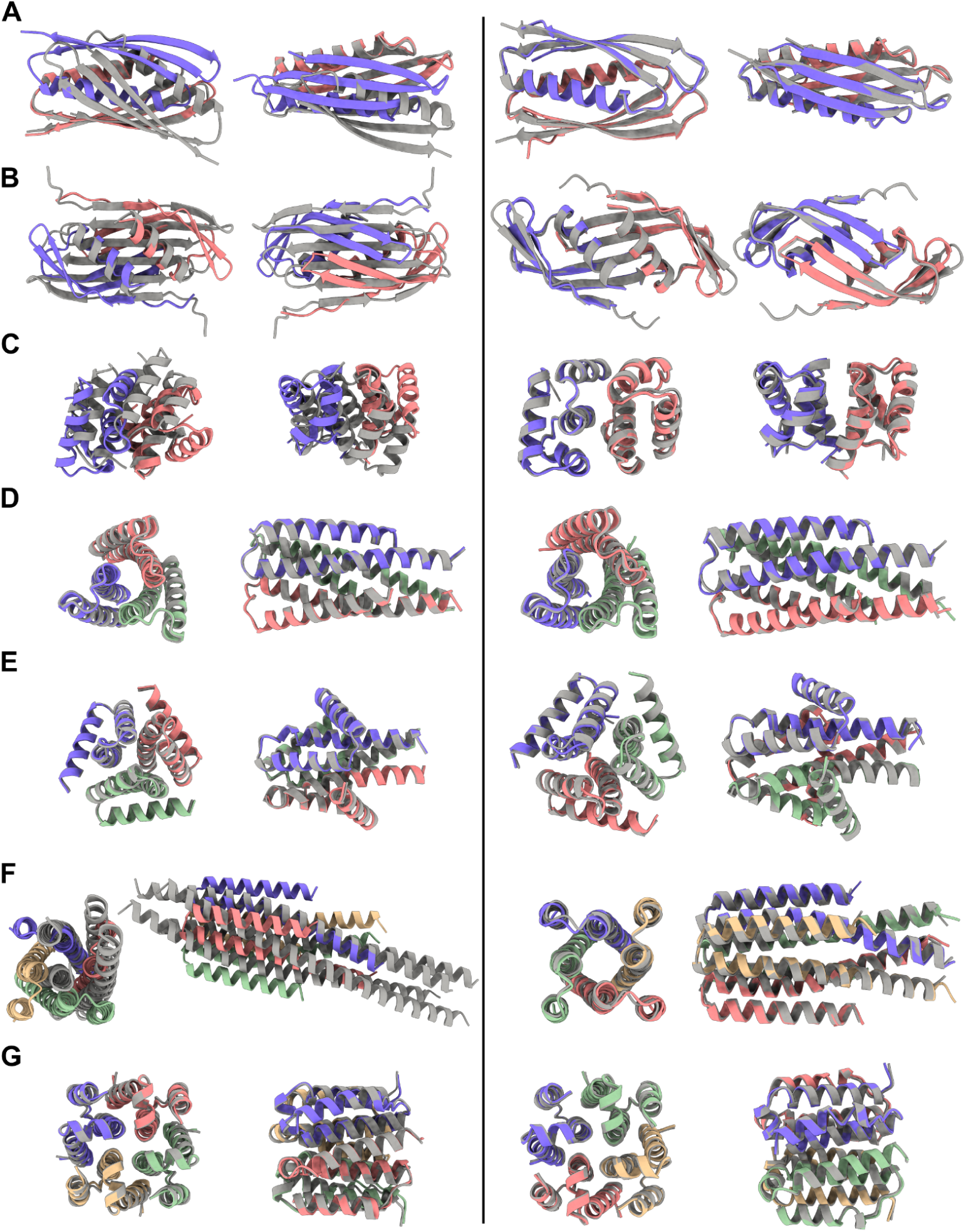
RoseTTAFold2 accurately predicts structures of crystallized HALs but not necessarily the original AF2 hallucinated backbone sequence. RoseTTAFold2 predictions compared to the original AF2 hallucination (left). RoseTTAFold2 prediction for the MPNN re-designs of the same backbones (right). (**A**) HALC2_062 (RMSD: 2.75 Å | 0.83 Å). (**B**) HALC2_065 (RMSD: 4.28 Å | 1.11 Å). (**C**) HALC2_068 (RMSD: 3.91 Å | 0.92 Å). (**D**) HALC3_104 (RMSD: 0.27 Å | 0.42 Å). (**E**) HALC3_109 (RMSD: 0.48 Å | 0.55 Å). (**F**) HALC4_135 (RMSD: 4.08 Å | 0.72 Å). (**G**) HALC4_136 (RMSD: 0.91 Å | 0.37 Å). The AF2/crystal structures are colored by chain, and the RoseTTAFold2 predictions are shown in gray.

**Fig. S7.**
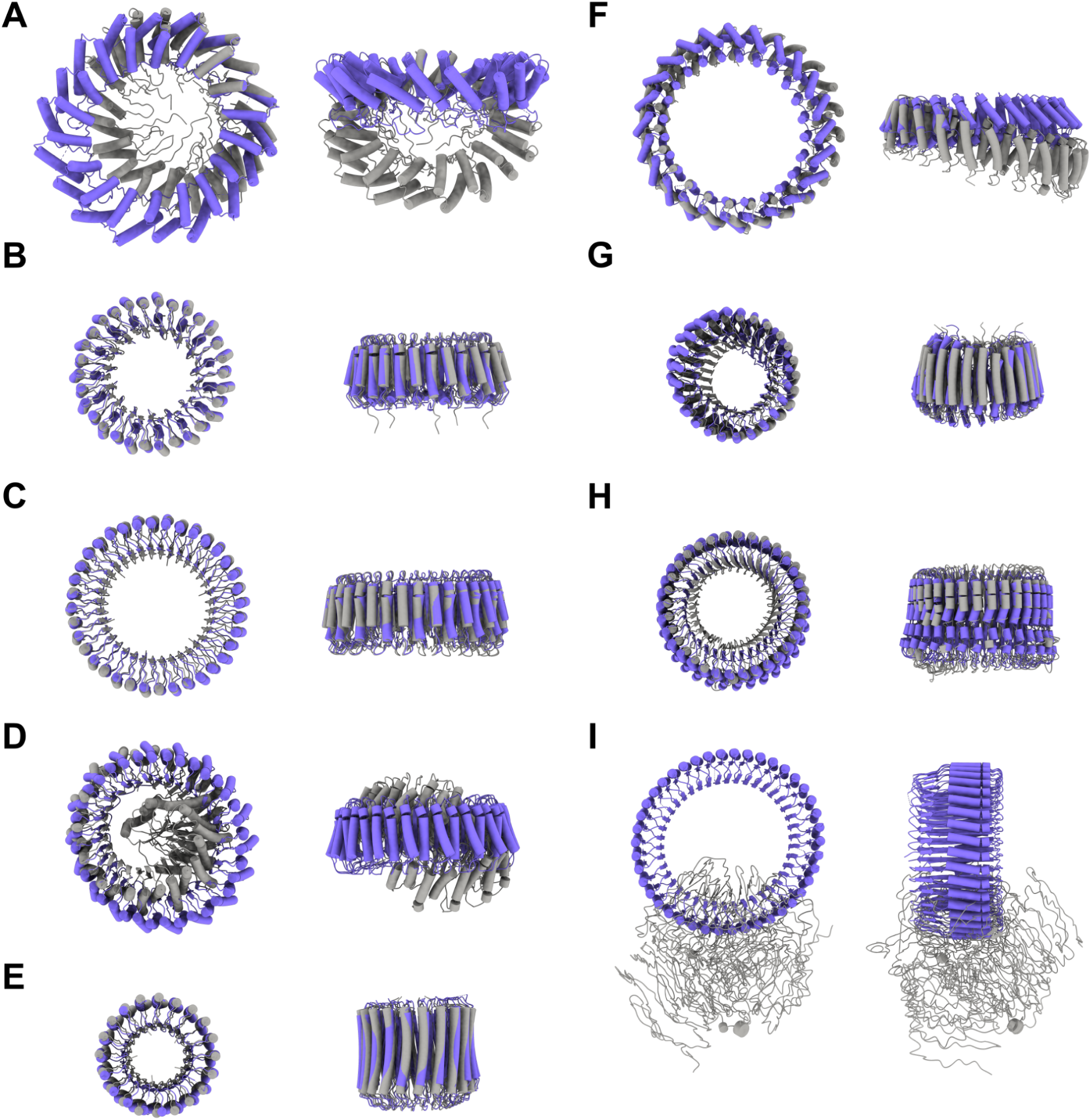
RoseTTAFold2 predictions aligned to design models of EM-verified HALs. Design ID (RMSD of AF2 model against RF2 prediction) (**A**) HALC6_220 (3.87 Å). (**B**) HALC15-5_262 (1.33 Å). (**C**) HALC18-6_265 (0.73 Å). (**D**) HALC18-6_278 (3.35 Å). (**E**) HALC20-5_308 (1.18 Å). (**F**) HALC24-6_316 (3.66 Å). (**G**) HALC25-5_341 (3.09 Å). (**H**) HALC33-3_343 (3.67 Å). (**I**) HALC42-7_351 (7.99 Å). RF2 predictions in gray, AF2 design models in purple.

**Fig. S8.**
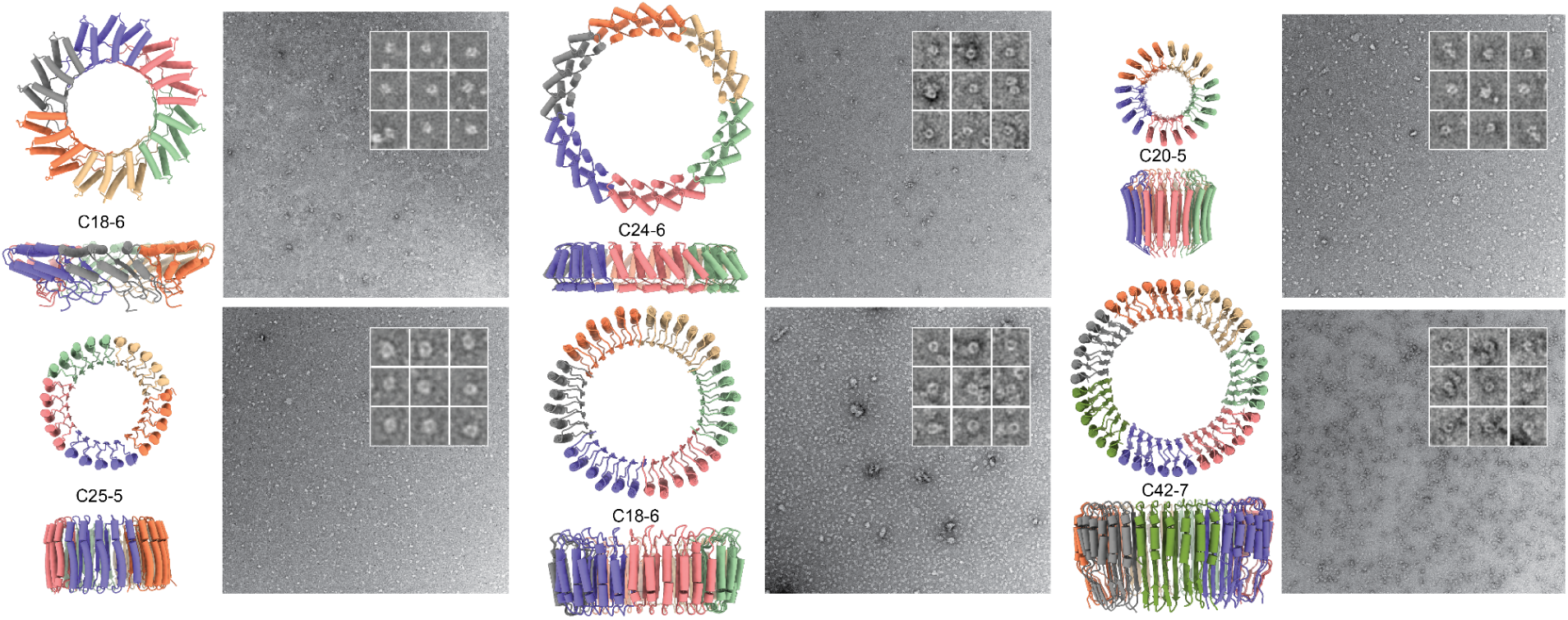
Design models and corresponding experimental negative stain electron microscopy analysis of designs shown in Fig. 3A. A raw micrograph at 57k magnification is shown along with nine example extracted particles that were used for further classification and data processing. From top left to bottom right: HALC6_220, HALC24-6_316, HALC20-5_308, HALC25-5_341, HALC18-6_278 and HALC42-7_351

**Fig. S9.**
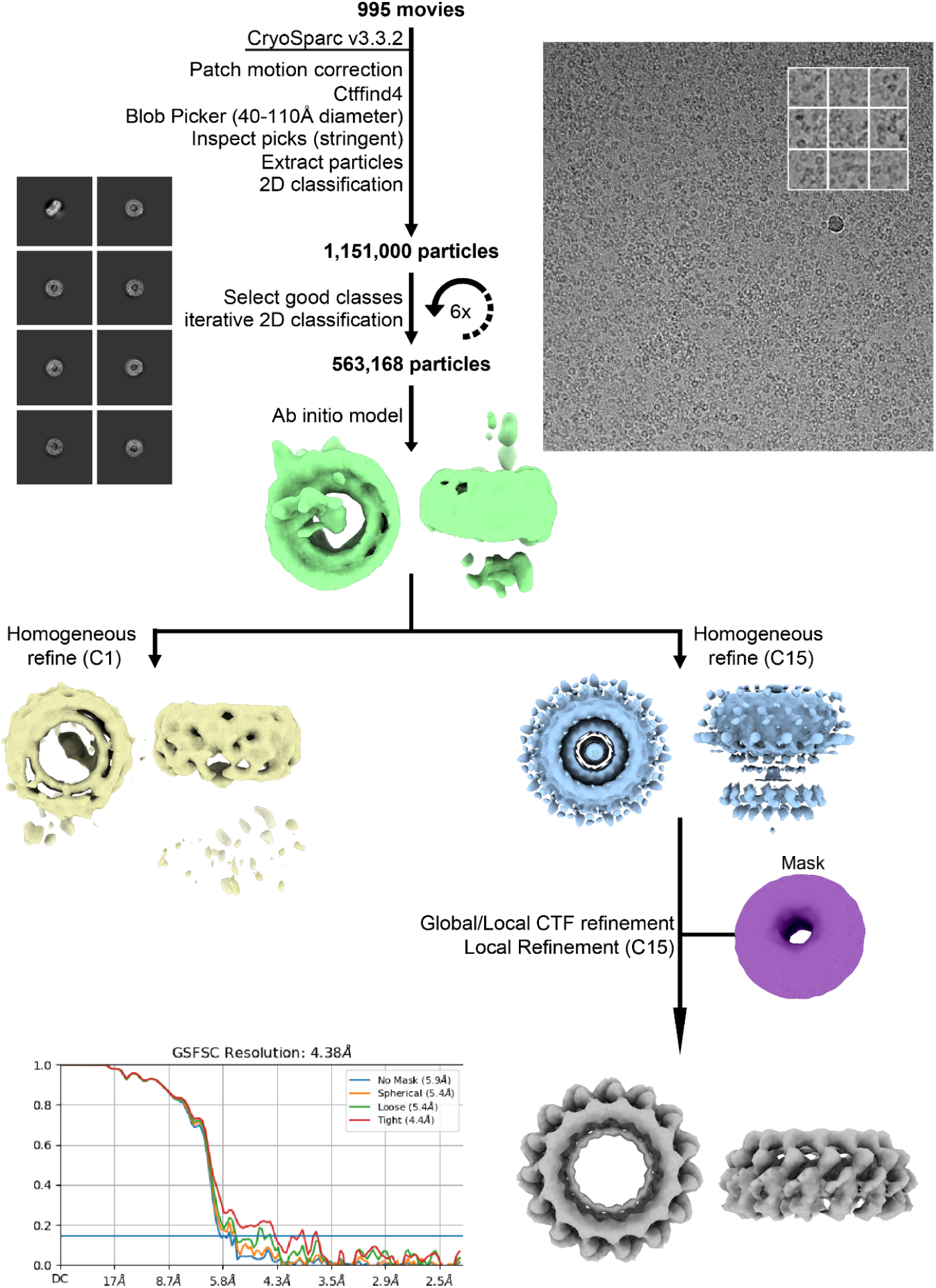
Details of cryoEM data processing pipeline used to generate electron density and structures of HALC15-5_262. 2D class averages and *ab initio* reconstruction followed by a C1 homogeneous refine yielded C15 features corresponding to the size and secondary structure of the design model, which allowed us to further process the design with C15 symmetry imposed here. A representative raw cryoEM micrograph is shown on the right along with nine example extracted particles and characteristic 2D class averages used in the processing pipeline. An FSC validation curve for the final reconstruction is shown along with the electron density map.

**Fig. S10.**
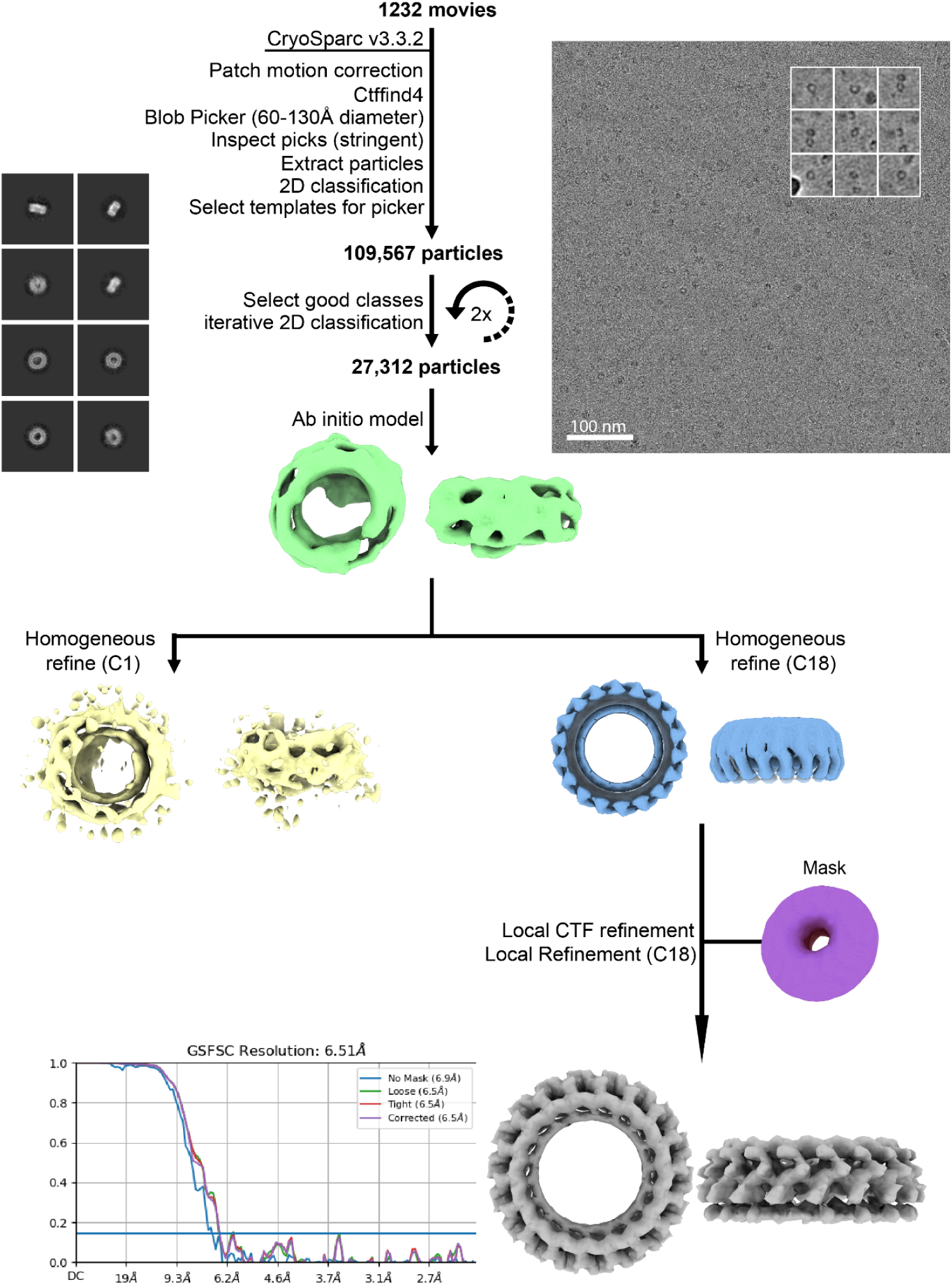
Details of cryoEM data processing pipeline used to generate electron density and structures of HALC18-6_265. 2D class averages and *ab initio* reconstruction followed by a C1 homogeneous refine yielded C18 features corresponding to the size and secondary structure of the design model, which allowed us to further process the design with C18 symmetry imposed here. A representative raw cryoEM micrograph is shown on the right along with nine example extracted particles and characteristic 2D class averages used in the processing pipeline. An FSC validation curve for the final reconstruction is shown along with the electron density map.

**FIG. S11.**
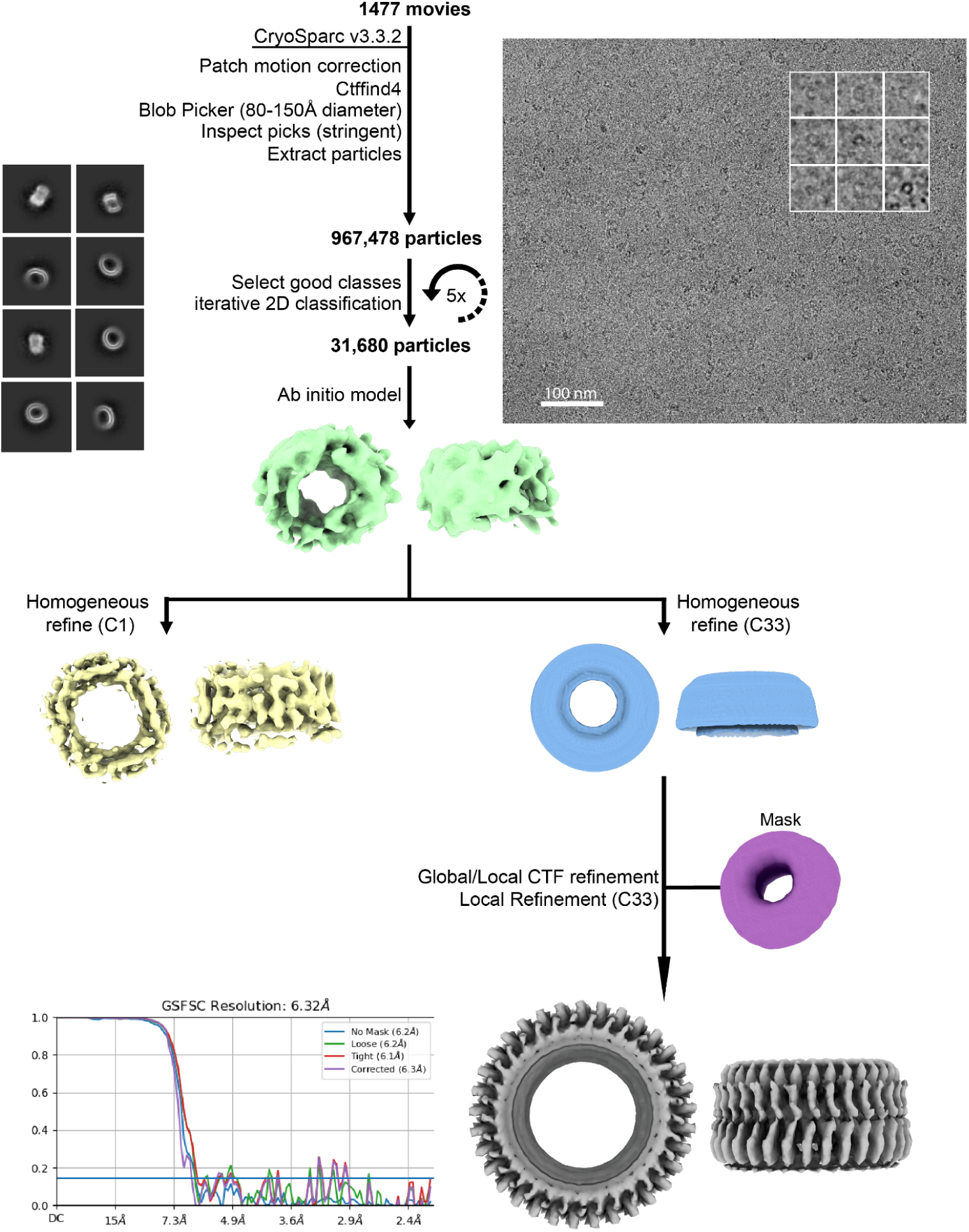
Details of cryoEM data processing pipeline used to generate electron density and structures of HALC33-3_343. 2D class averages and *ab initio* reconstruction followed by a C1 homogeneous refine yielded C33 features corresponding to the size and secondary structure of the design model, which allowed us to further process the design with C33 symmetry imposed here. A representative raw cryoEM micrograph is shown on the right along with nine example extracted particles and characteristic 2D class averages used in the processing pipeline. An FSC validation curve for the final reconstruction is shown along with the electron density map.

**FIG. S12.**
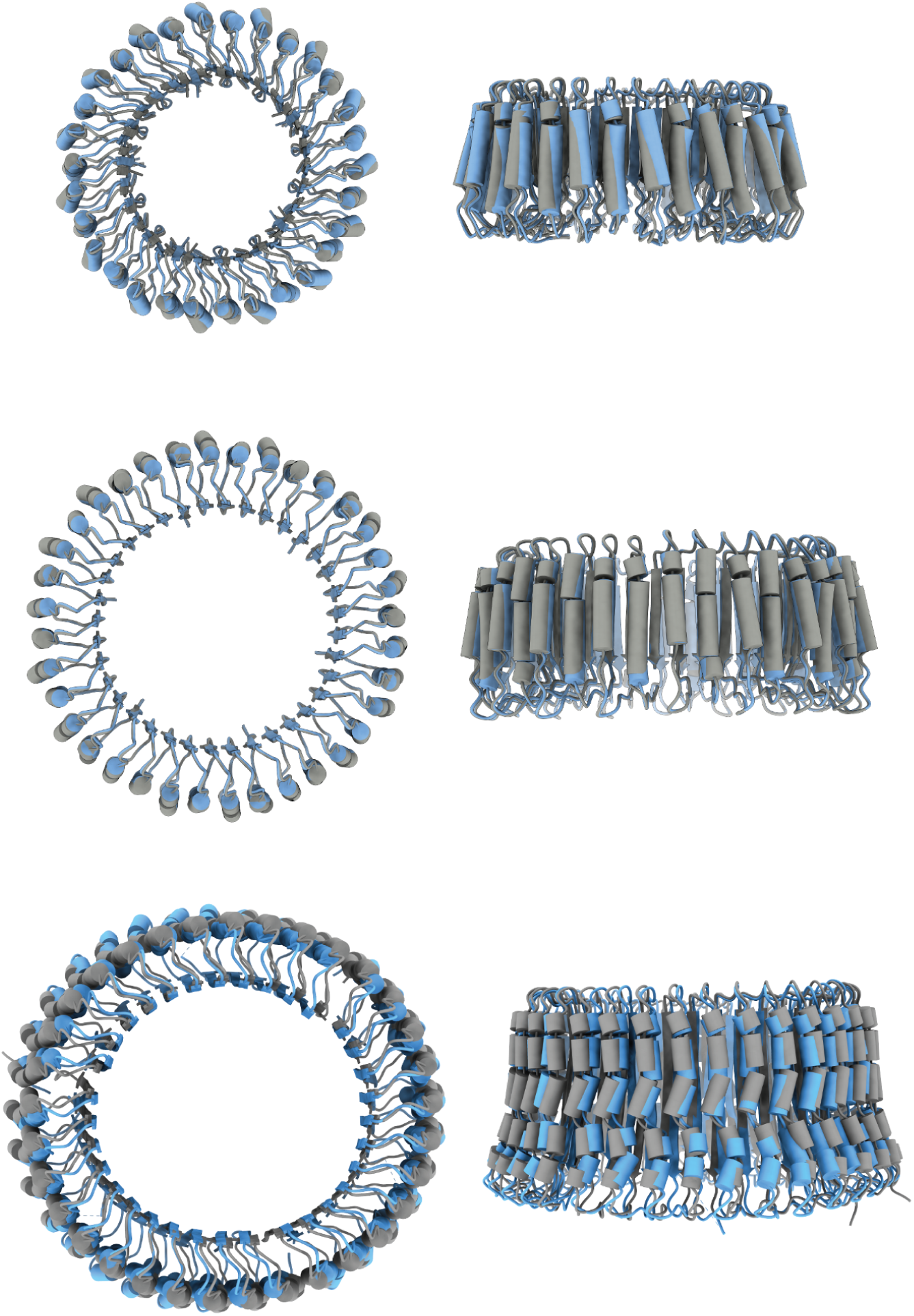
Detailed comparison of HAL designs versus cryoEM structures. The designs were relaxed into experimental cryoEM electron densities using Rosetta FastRelax and SetupForDensityScoring. From Top to Bottom: HALC15-5_262, HALC18-6_265, and HALC33-3_343. Superposition of the designed backbone (gray) and backbone relaxed into the experimental electron density (light blue). The computed backbone atom RMSD between the designed and experimental structure are 0.81 Å, 1.69 Å, and 2.30 Å respectively.

**FIG. S13.**
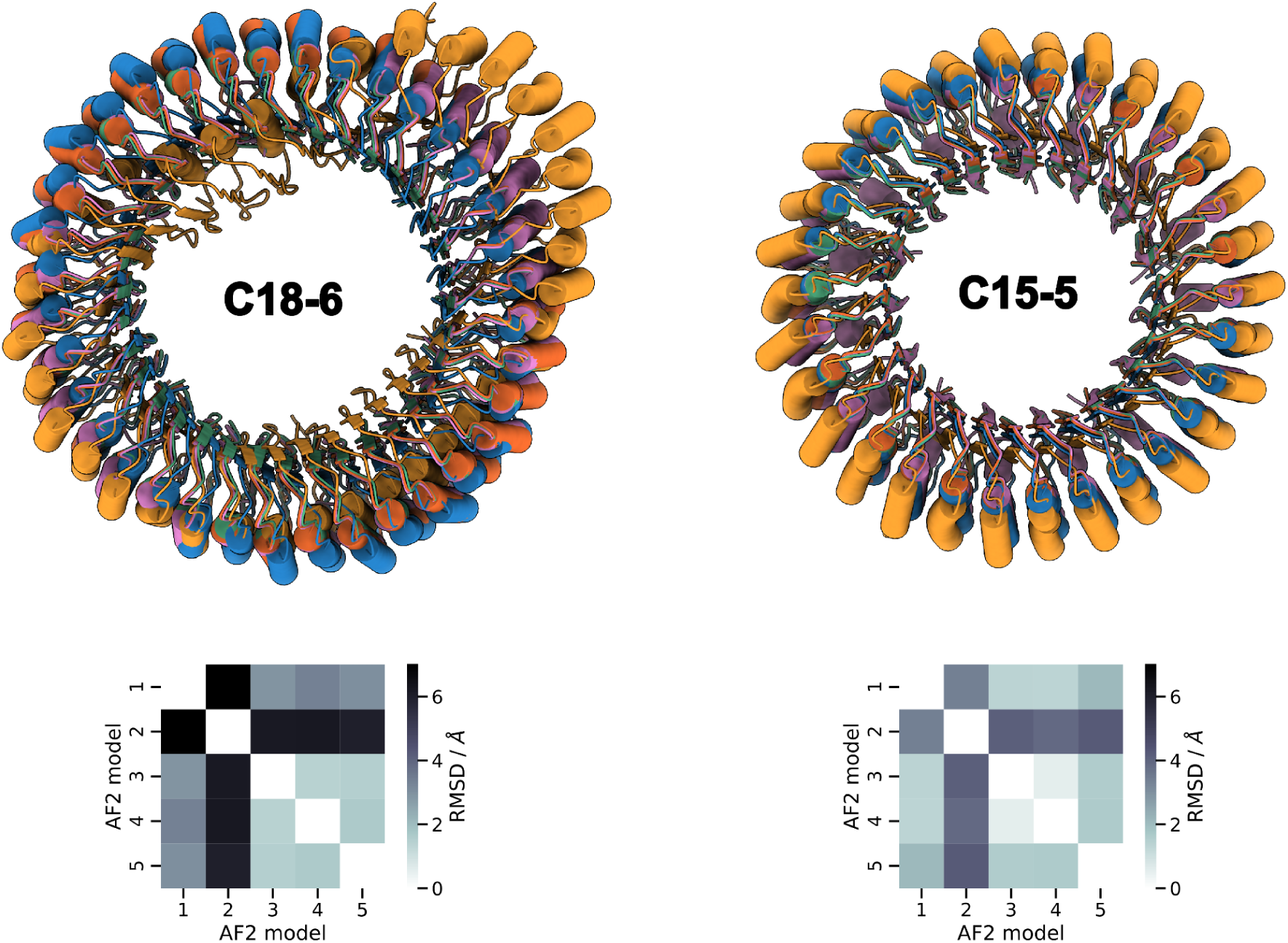
AF2 correctly predicts oligomeric valency. HALC15-5_262 was originally designed as a homo-hexamer but the cryoEM structure (Fig. 3B) revealed a homo-pentamer. Prediction of the sequence with all five models in both the homo-hexameric and homo-pentameric configurations reveals smaller structural deviations and higher confidence scores for the homo-pentamer. Mean values for homo-hexamer | homo-pentamer; pLDDT: 70.57 | 73.88, pTM: 0.486 | 0.57, pAE: 19.59 | 16.89. Top: structural alignments of all five models (model_1_ptm; blue, model_2_ptm; orange, model_3_ptm; green, model_4_ptm; red, model_5_ptm; pink). Bottom: all by all RMSD values between the five models.

**Table S1.**
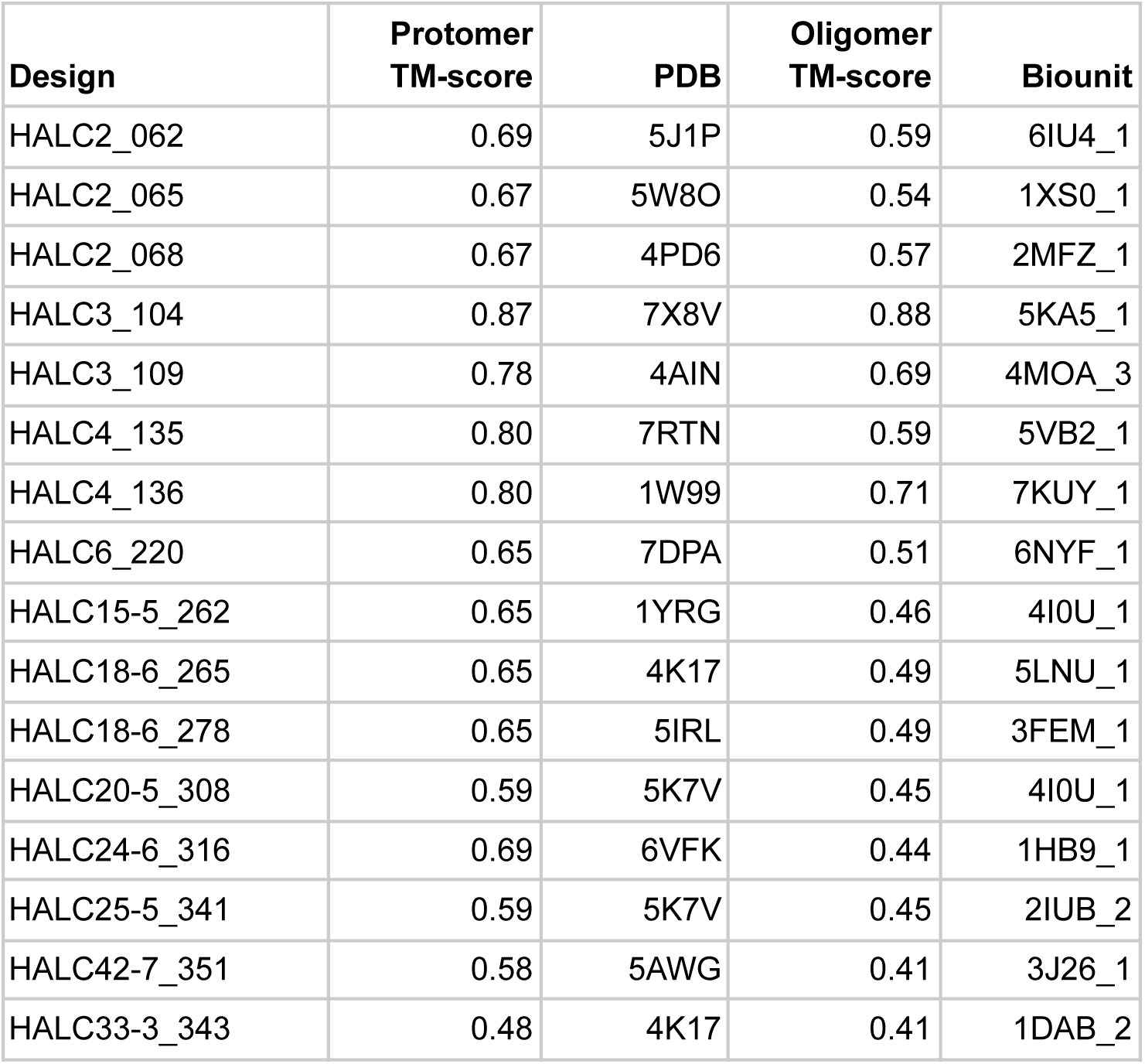
PDB IDs of the closest matches for structurally-validated HALs (Fig. 2-3).

| **Design** | **Protomer**<br>**TM-score** | **PDB** | **Oligomer**<br>**TM-score** | **Biounit** |
| --- | --- | --- | --- | --- |
| HALC2_062 | 0.69 | 5J1P | 0.59 | 6IU4_1 |
| HALC2_065 | 0.67 | 5W8O | 0.54 | 1XS0_1 |
| HALC2_068 | 0.67 | 4PD6 | 0.57 | 2MFZ_1 |
| HALC3_104 | 0.87 | 7X8V | 0.88 | 5KA5_1 |
| HALC3_109 | 0.78 | 4AIN | 0.69 | 4MOA_3 |
| HALC4_135 | 0.80 | 7RTN | 0.59 | 5VB2_1 |
| HALC4_136 | 0.80 | 1W99 | 0.71 | 7KUY_1 |
| HALC6_220 | 0.65 | 7DPA | 0.51 | 6NYF_1 |
| HALC15-5_262 | 0.65 | 1YRG | 0.46 | 4IOU_1 |
| HALC18-6_265 | 0.65 | 4K17 | 0.49 | 5LNU_1 |
| HALC18-6_278 | 0.65 | 5IRL | 0.49 | 3FEM_1 |
| HALC20-5_308 | 0.59 | 5K7V | 0.45 | 4IOU_1 |
| HALC24-6_316 | 0.69 | 6VFK | 0.44 | 1HB9_1 |
| HALC25-5_341 | 0.59 | 5K7V | 0.45 | 2IUB_2 |
| HALC42-7_351 | 0.58 | 5AWG | 0.41 | 3J26_1 |
| HALC33-3_343 | 0.48 | 4K17 | 0.41 | 1DAB_2 |

**Table S2.** UniRef100 IDs of the best hits for structurally-validated HALs (Fig. 2-3).

| **Design** | **Repeat**<br>**E-value** | **UniRef100 ID** | **Protomer**<br>**E-value** | **UniRef100 ID** |
| --- | --- | --- | --- | --- |
| HALC2_062 | 3.70E+00 | UPI00131BD06C | 3.70E+00 | UPI00131BD06C |
| HALC2_065 | 5.80E-01 | A0A8I1R8D5 | 5.80E-01 | A0A8I1R8D5 |
| HALC2_068 | 1.40E+00 | A0A6B2M0S8 | 1.40E+00 | A0A6B2M0S8 |
| HALC3_104 | 4.70E-01 | UPI0013B3A05C | 4.70E-01 | UPI0013B3A05C |
| HALC3_109 | 8.20E-01 | UPI000B0DAB1F | 8.20E-01 | UPI000B0DAB1F |
| HALC4_135 | 2.80E-01 | A0A3G1RPF3 | 2.80E-01 | A0A3G1RPF3 |
| HALC4_136 | 6.50E+00 | A7ANS2 | 6.50E+00 | A7ANS2 |
| HALC6_220 | 2.00E-02 | A0A434I672 | 2.00E-02 | A0A434I672 |
| HALC15-5_262 | 5.70E-02 | I7LU18 | 3.50E-17 | A0A7S2JY04 |
| HALC18-6_265 | 8.00E-03 | W2S5F8 | 3.17E-16 | A0A7S2JY04 |
| HALC18-6_278 | 5.00E-01 | A0A7E5WBQ0 | 2.99E-08 | A0A819R934 |
| HALC20-5_308 | 9.60E+00 | A0A1F4XIB2 | 1.13E-05 | UPI001CF37084 |
| HALC24-6_316 | 1.00E+01 | UPI0019D624AA | 3.00E-03 | A0A7G8BM39 |
| HALC25-5_341 | 2.60E+01 | A0A6N1YEJ1 | 1.86E-09 | A0A2B4S1A5 |
| HALC33-3_343 | 8.80E-01 | D7MIU3 | 1.62E-35 | A0A2I0HQ60 |
| HALC42-7_351 | 1.40E+01 | A0A7L1D0M5 | 1.35E-14 | B4SHG6 |

**Table S3.** Crystallographic statistics and PDB accession numbers for the structures displayed in Fig. 2.

|  | HALC2_062<br>PDB: 8D04 | HALC2_065<br>PDB: 8D03 | HALC2_068<br>PDB: 8D05 | HALC3_104<br>PDB: 8D06 | HALC3_109<br>PDB: 8D07 | HALC4_135<br>PDB: 8D08 | HALC4_136<br>PDB: 8D09 |
| --- | --- | --- | --- | --- | --- | --- | --- |
| Space group | P 65 | P 42 | P 32 2 1 | P 41 | C 1 2 1 | P 41 21 2 | C 2 2 21 |
| **Cell dimensions** |  |  |  |  |  |  |  |
| a, b, c (Å) | 67.9, 67.9,<br>228.4 | 50.2, 50.2,<br>22.1 | 70.6, 70.6,<br>31.4 | 107.5, 107.5,<br>111.7 | 136.8, 136.8,<br>94.2 | 35.9, 35.9,<br>438.0 | 52.8, 77.9,<br>52.8 |
| $\alpha, \beta, \gamma$ (°) | 90,90,120 | 90, 90, 90 | 90, 90, 120 | 90, 90, 90 | 90, 129.7, 90 | 90, 90, 90 | 90, 90, 90 |
| **Data Collection** |  |  |  |  |  |  |  |
| Resolution (Å)* | 56.95 - 2.11<br>(2.19 - 2.11) | 50.19 - 2.51<br>(2.60 - 2.51) | 20.39 - 1.75<br>(1.81 - 1.75) | 76.01 - 3.40<br>(3.52 - 3.40) | 72.61 - 2.09<br>(2.17 - 2.09) | 54.75 - 3.30<br>(3.41 - 3.30) | 23.62 - 1.90<br>(1.97 - 1.90) |
| Rmerge | 0.067 (2.197) | 0.311 (1.853) | 0.447 (1.368) | 0.076 (0.641) | 0.089 (0.684) | 0.148 (0.819) | 0.351 (0.884) |
| Rpim | 0.028 (0.878) | 0.089 (0.515) | 0.151 (0.496) | 0.037 (0.344) | 0.050 (0.400) | 0.060 (0.328) | 0.102 (0.244) |
| Mean I/$\sigma$(I) | 16.65 (1.17) | 2.85 (0.66) | 8.85 (1.33) | 14.56 (2.61) | 11.77 (2.06) | 9.29 (1.37) | 9.45 (3.95) |
| CC 1/2 | 0.996 (0.559) | 0.987 (0.336) | 0.95 (0.566) | 0.999 (0.748) | 0.995 (0.621) | 0.996 (0.639) | 0.981 (0.832) |
| Completeness (%) | 99.81 (99.47) | 99.90 (100) | 98.89 (89.99) | 99.40 (99.43) | 98.51 (99.27) | 98.67 (95.19) | 99.52 (100) |
| Redundancy | 7.1 (7.2) | 13.2 (14.0) | 9.8 (8.1) | 4.8 (4.3) | 3.8 (3.9) | 6.8 (6.4) | 13.1 (13.6) |
| **Refinement** |  |  |  |  |  |  |  |
| No. unique reflections | 34088 (3405) | 2002 (193) | 9287 (819) | 17541 (1749) | 20957 (2047) | 8009 (435) | 8869 (875) |
| Rwork / Rfree (%) | 23.6 (32.1) /<br>26.3 (33.4) | 24.2 (41.3) /<br>26.5 (34.8) | 19.0 (27.4) /<br>20.5 (26.2) | 28.4 (35.6) /<br>30.9 (38.3) | 20.6 (27.0) /<br>26.7 (34.8) | 25.0 (39.8) /<br>29.8 (43.7) | 23.2 (28.2) /<br>25.5 (30.5) |
| No. non-hydrogen atoms | 3210 | 469 | 563 | 6344 | 3159 | 2208 | 1056 |
| Macromolecules | 3210 | 469 | 538 | 6344 | 3159 | 2208 | 1020 |
| Solvent | 0 | 0 | 25 | 0 | 0 | 0 | 36 |
| Ramachandran favoured / allowed (%) | 96.52 / 3.48 | 94.83 / 5.17 | 98.41 / 1.59 | 97.33 / 2.67 | 99.20 / 0.80 | 92.58 / 7.42 | 99.21 / 0.79 |
| **R.m.s. deviations** |  |  |  |  |  |  |  |
| Bond lengths (Å) | 0.003 | 0.002 | 0.006 | 0.003 | 0.007 | 0.011 | 0.014 |
| Bond angles (°) | 0.51 | 0.48 | 0.77 | 0.53 | 0.92 | 1.37 | 1.47 |
| **B-factors (Å<sup>2</sup>)** |  |  |  |  |  |  |  |
| Macromolecules | 76.64 | 74.11 | 35.68 | 139.29 | 54.51 | 134.25 | 37.12 |
| Solvent |  |  | 42.86 |  |  |  | 44.17 |

\*Statistics for the highest-resolution shell are shown in parentheses

## References

1. H. Garcia-Seisdedos, C. Empereur-Mot, N. Elad, E. D. Levy, Proteins evolve on the edge of supramolecular self-assembly. Nature. 548, 244–247 (2017).

2. I. G. Johnston, K. Dingle, S. F. Greenbury, C. Q. Camargo, J. P. K. Doye, S. E. Ahnert, A. A. Louis, Symmetry and simplicity spontaneously emerge from the algorithmic nature of evolution. Proc. Natl. Acad. Sci. 119, e2113883119 (2022).

3. S. E. Ahnert, J. A. Marsh, H. Hernández, C. V. Robinson, S. A. Teichmann, Principles of assembly reveal a periodic table of protein complexes. Science. 350, aaa2245 (2015).

4. wwPDB consortium, Protein Data Bank: the single global archive for 3D macromolecular structure data. Nucleic Acids Res. 47, D520–D528 (2019).

5. D. S. Goodsell, A. J. Olson, Structural Symmetry and Protein Function. Annu. Rev. Biophys. Biomol. Struct. 29, 105–153 (2000).

6. T. Handel, W. F. DeGrado, De novo design of a Zn2+-binding protein. J. Am. Chem. Soc. 112, 6710–6711 (1990).

7. P. B. Harbury, J. J. Plecs, B. Tidor, T. Alber, P. S. Kim, High-Resolution Protein Design with Backbone Freedom. Science. 282, 1462–1467 (1998).

8. J. A. Fallas, G. Ueda, W. Sheffler, V. Nguyen, D. E. McNamara, B. Sankaran, J. H. Pereira, F. Parmeggiani, T. J. Brunette, D. Cascio, T. R. Yeates, P. Zwart, D. Baker, Computational design of self-assembling cyclic protein homo-oligomers. Nat. Chem. 9, 353–360 (2017).

9. A. R. Thomson, C. W. Wood, A. J. Burton, G. J. Bartlett, R. B. Sessions, R. L. Brady, D. N. Woolfson, Computational design of water-soluble α-helical barrels. Science. 346, 485–488 (2014).

10. P.-S. Huang, K. Feldmeier, F. Parmeggiani, D. A. Fernandez Velasco, B. Höcker, D. Baker, De novo design of a four-fold symmetric TIM-barrel protein with atomic-level accuracy. Nat. Chem. Biol. 12, 29–34 (2016).

11. P.-S. Huang, G. Oberdorfer, C. Xu, X. Y. Pei, B. L. Nannenga, J. M. Rogers, F. DiMaio, T. Gonen, B. Luisi, D. Baker, High thermodynamic stability of parametrically designed helical bundles. Science. 346, 481–485 (2014).

12. S. E. Boyken, Z. Chen, B. Groves, R. A. Langan, G. Oberdorfer, A. Ford, J. M. Gilmore, C. Xu, F. DiMaio, J. H. Pereira, B. Sankaran, G. Seelig, P. H. Zwart, D. Baker, De novo design of protein homo-oligomers with modular hydrogen-bond network–mediated specificity. Science. 352, 680–687 (2016).

13. J. B. Bale, S. Gonen, Y. Liu, W. Sheffler, D. Ellis, C. Thomas, D. Cascio, T. O. Yeates, T. Gonen, N. P. King, D. Baker, Accurate design of megadalton-scale two-component icosahedral protein complexes. Science. 353, 389–394 (2016).

14. I. Vulovic, Q. Yao, Y.-J. Park, A. Courbet, A. Norris, F. Busch, A. Sahasrabuddhe, H. Merten, D. D. Sahtoe, G. Ueda, J. A. Fallas, S. J. Weaver, Y. Hsia, R. A. Langan, A. Plückthun, V. H. Wysocki, D. Veesler, G. J. Jensen, D. Baker, Generation of ordered protein assemblies using rigid three-body fusion. Proc. Natl. Acad. Sci. 118, e2015037118 (2021).

15. Y. Hsia, R. Mout, W. Sheffler, N. I. Edman, I. Vulovic, Y.-J. Park, R. L. Redler, M. J. Bick, A. K. Bera, A. Courbet, A. Kang, T. J. Brunette, U. Nattermann, E. Tsai, A. Saleem, C. M. Chow, D. Ekiert, G. Bhabha, D. Veesler, D. Baker, Design of multi-scale protein complexes by hierarchical building block fusion. Nat. Commun. 12, 2294 (2021).

16. C. E. Correnti, J. P. Hallinan, L. A. Doyle, R. O. Ruff, C. A. Jaeger-Ruckstuhl, Y. Xu, B. W. Shen, A. Qu, C. Polkinghorn, D. J. Friend, A. D. Bandaranayake, S. R. Riddell, B. K. Kaiser, B. L. Stoddard, P. Bradley, Engineering and functionalization of large circular tandem repeat protein nanoparticles. Nat. Struct. Mol. Biol. 27, 342–350 (2020).

17. D. D. Sahtoe, F. Praetorius, A. Courbet, Y. Hsia, B. I. M. Wicky, N. I. Edman, L. M. Miller, B. J. R. Timmermans, J. Decarreau, H. M. Morris, A. Kang, A. K. Bera, D. Baker, Reconfigurable asymmetric protein assemblies through implicit negative design. Science. 375, eabj7662 (2022).

18. I. Anishchenko, S. J. Pellock, T. M. Chidyausiku, T. A. Ramelot, S. Ovchinnikov, J. Hao, K. Bafna, C. Norn, A. Kang, A. K. Bera, F. DiMaio, L. Carter, C. M. Chow, G. T. Montelione, D. Baker, De novo protein design by deep network hallucination. Nature. 600, 547–552 (2021).

19. M. Jendrusch, J. O. Korbel, S. K. Sadiq, AlphaDesign: A de novo protein design framework based on AlphaFold (2021), p. 2021.10.11.463937, doi:10.1101/2021.10.11.463937.

20. L. Moffat, J. G. Greener, D. T. Jones, Using AlphaFold for Rapid and Accurate Fixed Backbone Protein Design (2021), p. 2021.08.24.457549, doi:10.1101/2021.08.24.457549.

21. J. Wang, S. Lisanza, D. Juergens, D. Tischer, I. Anishchenko, M. Baek, J. L. Watson, J. H. Chun, L. F. Milles, J. Dauparas, M. Expòsit, W. Yang, A. Saragovi, S. Ovchinnikov, D. Baker, Deep learning methods for designing proteins scaffolding functional sites (2021), p. 2021.11.10.468128, doi:10.1101/2021.11.10.468128.

22. S. Ovchinnikov, P.-S. Huang, Structure-based protein design with deep learning. Curr. Opin. Chem. Biol. 65, 136–144 (2021).

23. C. Norn, B. I. M. Wicky, D. Juergens, S. Liu, D. Kim, D. Tischer, B. Koepnick, I. Anishchenko, Foldit Players, D. Baker, S. Ovchinnikov, A. Coral, A. J. Bubar, A. Boykov, A. U. Valle Pérez, A. MacMillan, A. Lubow, A. Mussini, A. Cai, A. J. Ardill, A. Seal, A. Kalantarian, B. Failer, B. Lackersteen, B. Chagot, B. R. Haight, B. Taştan, B. Uitham, B. G. Roy, B. R. de Melo Cruz, B. Echols, B. E. Lorenz, B. Blair, B. Kestemont, C. D. Eastlake, C. J. Bragdon, C. Vardeman, C. Salerno, C. Comisky, C. L. Hayman, C. R. Landers, C. Zimov, C. D. Coleman, C. R. Painter, C. Ince, C. Lynagh, D. Malaniia, D. C. Wheeler, D. Robertson, V. Simon, E. Chisari, E. L. J. Kai, F. Rezae, F. Lengyel, F. Tabotta, F. Padelletti, F. Boström, G. O. Gross, G. McIlvaine, G. Beecher, G. T. Hansen, G. de Jong, H. Feldmann, J. L. Borman, J. Quinn, J. Norrgard, J. Truong, J. A. Diderich, J. M. Canfield, J. Photakis, J. D. Slone, J. Madzio, J. Mitchell, J. C. Stomieroski, J. H. Mitch, J. R. Altenbeck, J. Schinkler, J. B. Weinberg, J. D. Burbach, J. C. Sequeira da Costa, J. F. Bada Juarez, J. P. Gunnarsson, K. D. Harper, K. Joo, K. T. Clayton, K. E. DeFord, K. F. Scully, K. M. Gildea, K. J. Abbey, K. L. Kohli, K. Stenner, K. Takács, L. L. Poussaint, L. C. Manalo, L. C. Withers, L. Carlson, L. Wei, L. R. Fisher, L. Carpenter, M. Ji-hwan, M. Ricci, M. A. Belcastro, M. Leniec, M. Hohmann, M. Thompson, M. A. Thayer, M. Gaebel, M. D. Cassidy, M. Fagiola, M. Lewis, M. Pfützenreuter, M. Simon, M. M. Elmassry, N. Benevides, N. K. Kerr, N. Verma, O. Shannon, O. Yin, P. Wolfteich, P. Gummersall, P. Tłuścik, P. Gajar, P. J. Triggiani, R. Guha, R. B. Mathew Innes, R. Buchanan, R. Gamble, R. Leduc, R. Spearing, R. L. C. dos Santos Gomes, R. D. Estep, R. DeWitt, R. Moore, S. G. Shnider, S. J. Zaccanelli, S. Kuznetsov, S. Burillo-Sanz, S. Mooney, S. Vasiliy, S. S. Butkovich, S. B. Hudson, S. L. Pote, S. P. Denne, S. A. Schwegmann, S. Ratna, S. C. Kleinfelter, T. Bausewein, T. J. George, T. S. de Almeida, U. Yeginer, W. Barmettler, W. R. Pulley, W. S. Wright, Willyanto, W. Lansford, X. Hochart, Y. A. S. Gaiji, Y. Lagodich, V. Christian, Protein sequence design by conformational landscape optimization. Proc. Natl. Acad. Sci. 118, e2017228118 (2021).

24. N. Anand, R. Eguchi, I. I. Mathews, C. P. Perez, A. Derry, R. B. Altman, P.-S. Huang, Protein sequence design with a learned potential. Nat. Commun. 13, 746 (2022).

25. J. Jumper, R. Evans, A. Pritzel, T. Green, M. Figurnov, O. Ronneberger, K. Tunyasuvunakool, R. Bates, A. Žídek, A. Potapenko, A. Bridgland, C. Meyer, S. A. A. Kohl, A. J. Ballard, A. Cowie, B. Romera-Paredes, S. Nikolov, R. Jain, J. Adler, T. Back, S. Petersen, D. Reiman, E. Clancy, M. Zielinski, M. Steinegger, M. Pacholska, T. Berghammer, S. Bodenstein, D. Silver, O. Vinyals, A. W. Senior, K. Kavukcuoglu, P. Kohli, D. Hassabis, Highly accurate protein structure prediction with AlphaFold. Nature. 596, 583–589 (2021).

26. J. Xu, Y. Zhang, How significant is a protein structure similarity with TM-score = 0.5? Bioinformatics. 26, 889–895 (2010).

27. Inceptionism: Going Deeper into Neural Networks. Google AI Blog, (available at http://ai.googleblog.com/2015/06/inceptionism-going-deeper-into-neural.html).

28. A. Nguyen, J. Yosinski, J. Clune, Deep Neural Networks are Easily Fooled: High Confidence Predictions for Unrecognizable Images (2015), (available at http://arxiv.org/abs/1412.1897).

29. K. Simonyan, A. Vedaldi, A. Zisserman, Deep Inside Convolutional Networks: Visualising Image Classification Models and Saliency Maps (2014), (available at http://arxiv.org/abs/1312.6034).

30. M. Baek, F. DiMaio, I. Anishchenko, J. Dauparas, S. Ovchinnikov, G. R. Lee, J. Wang, Q. Cong, L. N. Kinch, R. D. Schaeffer, C. Millán, H. Park, C. Adams, C. R. Glassman, A. DeGiovanni, J. H. Pereira, A. V. Rodrigues, A. A. van Dijk, A. C. Ebrecht, D. J. Opperman, T. Sagmeister, C. Buhlheller, T. Pavkov-Keller, M. K. Rathinaswamy, U. Dalwadi, C. K. Yip, J. E. Burke, K. C. Garcia, N. V. Grishin, P. D. Adams, R. J. Read, D. Baker, Accurate prediction of protein structures and interactions using a three-track neural network. Science. 373, 871–876 (2021).

31. B. Kobe, J. Deisenhofer, The leucine-rich repeat: a versatile binding motif. Trends Biochem. Sci. 19, 415–421 (1994).

32. P. Guerra, M. González-Alamos, A. Llauró, A. Casañas, J. Querol-Audí, P. J. de Pablo, N. Verdaguer, Symmetry disruption commits vault particles to disassembly. Sci. Adv. 8, eabj7795 (2022).

33. A. Courbet, J. Hansen, Y. Hsia, N. Bethel, Y.-J. Park, C. Xu, A. Moyer, S. E. Boyken, G. Ueda, U. Nattermann, D. Nagarajan, D.-A. Silva, W. Sheffler, J. Quispe, A. Nord, N. King, P. Bradley, D. Veesler, J. Kollman, D. Baker, Computational design of mechanically coupled axle-rotor protein assemblies. Science. 376, 383–390 (2022).

## Supplementary References

34. Y. Zhang, J. Skolnick, TM-align: a protein structure alignment algorithm based on the TM-score. Nucleic Acids Res. 33, 2302–2309 (2005).

35. S. Mukherjee, Y. Zhang, MM-align: a quick algorithm for aligning multiple-chain protein complex structures using iterative dynamic programming. Nucleic Acids Res. 37, e83 (2009).

36. B. Dang, M. Mravic, H. Hu, N. Schmidt, B. Mensa, W. F. DeGrado, SNAC-tag for sequence-specific chemical protein cleavage. Nat. Methods. 16, 319–322 (2019).

37. W. Kabsch, XDS. Acta Crystallogr. D Biol. Crystallogr. 66, 125–132 (2010).

38. M. D. Winn, C. C. Ballard, K. D. Cowtan, E. J. Dodson, P. Emsley, P. R. Evans, R. M. Keegan, E. B. Krissinel, A. G. W. Leslie, A. McCoy, S. J. McNicholas, G. N. Murshudov, N. S. Pannu, E. A. Potterton, H. R. Powell, R. J. Read, A. Vagin, K. S. Wilson, Overview of the CCP4 suite and current developments. Acta Crystallogr. D Biol. Crystallogr. 67, 235–242 (2011).

39. A. J. McCoy, R. W. Grosse-Kunstleve, P. D. Adams, M. D. Winn, L. C. Storoni, R. J. Read, Phaser crystallographic software. J. Appl. Crystallogr. 40, 658–674 (2007).

40. P. Emsley, K. Cowtan, Coot: model-building tools for molecular graphics. Acta Crystallogr. D Biol. Crystallogr. 60, 2126–2132 (2004).

41. P. D. Adams, P. V. Afonine, G. Bunkóczi, V. B. Chen, I. W. Davis, N. Echols, J. J. Headd, L.-W. Hung, G. J. Kapral, R. W. Grosse-Kunstleve, A. J. McCoy, N. W. Moriarty, R. Oeffner, R. J. Read, D. C. Richardson, J. S. Richardson, T. C. Terwilliger, P. H. Zwart, PHENIX: a comprehensive Python-based system for macromolecular structure solution. Acta Crystallogr. D Biol. Crystallogr. 66, 213–221 (2010).

42. G. N. Murshudov, A. A. Vagin, E. J. Dodson, Refinement of Macromolecular Structures by the Maximum-Likelihood Method. Acta Crystallogr. D Biol. Crystallogr. 53, 240–255 (1997).

43. C. J. Williams, J. J. Headd, N. W. Moriarty, M. G. Prisant, L. L. Videau, L. N. Deis, V. Verma, D. A. Keedy, B. J. Hintze, V. B. Chen, S. Jain, S. M. Lewis, W. B. Arendall III, J. Snoeyink, P. D. Adams, S. C. Lovell, J. S. Richardson, D. C. Richardson, MolProbity: More and better reference data for improved all-atom structure validation. Protein Sci. 27, 293–315 (2018).

44. B. L. Nannenga, M. G. Iadanza, B. S. Vollmar, T. Gonen, Curr. Protoc. Protein Sci., in press, doi:10.1002/0471140864.ps1715s72.

45. T. Grant, A. Rohou, N. Grigorieff, cisTEM, user-friendly software for single-particle image processing. eLife. 7, e35383 (2018).

46. A. Punjani, J. L. Rubinstein, D. J. Fleet, M. A. Brubaker, cryoSPARC: algorithms for rapid unsupervised cryo-EM structure determination. Nat. Methods. 14, 290–296 (2017).

47. A. Punjani, D. J. Fleet, 3D variability analysis: Resolving continuous flexibility and discrete heterogeneity from single particle cryo-EM. J. Struct. Biol. 213, 107702 (2021).

48. B. Carragher, N. Kisseberth, D. Kriegman, R. A. Milligan, C. S. Potter, J. Pulokas, A. Reilein, Leginon: An Automated System for Acquisition of Images from Vitreous Ice Specimens. J. Struct. Biol. 132, 33–45 (2000).

49. S. Q. Zheng, E. Palovcak, J.-P. Armache, K. A. Verba, Y. Cheng, D. A. Agard, MotionCor2: anisotropic correction of beam-induced motion for improved cryo-electron microscopy. Nat. Methods. 14, 331–332 (2017).

50. A. Rohou, N. Grigorieff, CTFFIND4: Fast and accurate defocus estimation from electron micrographs. J. Struct. Biol. 192, 216–221 (2015).

